# Optimizing genome assembly, chromosome synteny, and genetic variant discovery from Oxford Nanopore sequences of *Dermacentor reticulatus* ticks

**DOI:** 10.64898/2026.09.18.752137

**Authors:** Katie C. Dillon, Hein Sprong, Isobel Ronai, Saurav Choudhary, Courtney Grant, Rodrigo de Paula Baptista, David A. Ray, Travis C. Glenn

## Abstract

Generating high-quality genome assemblies for small animals with large genomes is complex due to their small body size, DNA contamination, and repetitive elements. Ticks exemplify these complexities, while also being a global health threat to humans, domestic animals, and wildlife. Advances in long-read sequencing platforms now make it feasible to obtain large amounts of raw sequence data from individual specimens, but challenges remain. Key genome workflow challenges include error correction, assembly, transposable element annotation, and chromosome assignment and synteny. Here we examine three individual *Dermacentor reticulatus* ticks using deep Oxford Nanopore sequencing. Comparing and contrasting bioinformatic tools for raw read and assembly manipulation allows us to identify the parameters that provide a high-quality haploid genome assembly among 38 assemblies. We find Dorado error corrected raw read data from approximately two flow cells are needed, but limiting assembly coverage to 40x produces the highest quality assemblies. Then we examined the effects of optimal workflow parameters on downstream analyses of gene synteny and manual transposable element annotation of the three tick assemblies in comparison to an independent assembly of a fourth *D. reticulatus* individual from a different country. Gene synteny analysis allows chromosome assignment of scaffolds and transposable element identification was improved markedly with manual curation. Finally, we compared the genetic variation of *D. reticulatus* across two populations and found similar genetic diversity. Our study provides a clear workflow to obtain high-quality assemblies from Oxford Nanopore sequences and genetic characterization of a species with a large, complex, and repetitive genome.

## INTRODUCTION

Anthropogenic disturbances to the environment are driving major changes in infectious disease risk (Baker et al., 2022), including tick-borne diseases. Ticks are increasing due to urbanization (Diuk-Wasser, Fernandez, & Vanwambeke, 2025) and climate change is having a major impact on tick populations (Gilbert, 2021). The tick species *Dermacentor reticulatus* is abundant in Europe and western Russia (Földvári, Široký, Szekeres, Majoros, & Sprong, 2016; Rubel et al., 2016) where urbanization has yielded an increase in tick density in green spaces within urban areas, outnumbering *Ixodes ricinus* ticks in some areas of Europe (Olivieri, Gazzonis, Zanzani, Veronesi, & Manfredi, 2017; Zając, Woźniak, & Kulisz, 2026). *D. reticulatus* is tolerant to changing climates (Földvári et al., 2016; Zając, Bartosik, Kulisz, & Woźniak, 2020), and is predicted to experience population expansion into southern Europe (Noll et al., 2023). As ecological disturbances are increasing and *D. reticulatus* is the primary vector of medically (e.g., Omsk hemorrhagic fever) and veterinary important diseases (e.g., canine babesiosis, equine piroplasmosis) (Földvári et al., 2016), tick control strategies are needed.

Tick control strategies have traditionally relied on cell culture, attenuation, animal testing, and trial-and-error (George, Pound, & Davey, 2004; Sette & Rappuoli, 2010). These traditional approaches to developing tick control strategies can be time consuming, costly, and laborious (Ghosh, Azhahianambi, & Yadav, 2007). In contrast, approaches that leverage DNA sequence information increases the efficiency and robustness of anti-tick drugs, vaccines, and acaricides (Abbas, Jmel, Mekki, Dijkgraaf, & Kotsyfakis, 2023; Sette & Rappuoli, 2010). Only ∼2.1% of tick species currently have a genome assembly available (Dillon, Frederick, Sprong, Glenn, & Ronai, 2026). Therefore, additional tick genome assemblies and genome-wide assessments of genetic variation are needed to improve and develop new tick control strategies.

High-quality genome assemblies for ticks have been increasingly generated over the past six years due to the establishment of long-read third-generation sequencing by Oxford Nanopore Technologies (ONT) and Pacific Biosciences (PacBio) (Dillon et al., 2026). ONT is often the sequencing platform of choice for molecular ecology studies (Mikheyev & Tin, 2014) because it generates long reads with lower input DNA requirements, uses lower-cost instruments, and has a lower barrier to entry when compared to Illumina and PacBio (Yunhao Wang, Zhao, Bollas, Wang, & Au, 2021). However, best practices for using ONT are needed to yield high-quality genome assemblies and characterization. Additionally, there are few examples of the practical impact of improved ONT sequencing reagents (e.g., R9 vs. R10) and software (e.g., Guppy v4 vs. v6) on large complex genome assemblies.

ONT reads can capture the complexity of repetitive sequences in eukaryotic genomes (Shahid & Slotkin, 2020). Ticks are an ideal case study for characterizing transposable elements (TEs) as they have complex genomes with up to 69% being TEs (Ronai et al., 2026). Typically, TE curation relies on automated bioinformatic tools, but manual TE libraries provide the most accurate representation of the repetitive makeup of the species, as well as a resource for comparative genomics of ticks (Dillon et al., 2026).

Here we use ONT sequencing on three individual *D. reticulatus* ticks conducted with R9 reagents and then repeated with R10 reagents. First, we compare the impact of sequencing and bioinformatics (flow cells, base-calling, and assembly parameters) on genome assembly quality to provide guidelines for highly repetitive genomes. Second, we conduct automated gene annotation, manual TE curation, synteny analyses, and variant analyses to gain a deeper understanding of the *D. reticulatus* genome. These genome guidelines and resources help provide the foundation for future work on tick pangenomes. Finally, although a few species of ticks have assemblies available for multiple individuals (Baede et al., 2024; Forth et al., 2020; Ronai et al., 2026), significant questions remain regarding the variation among individuals within tick species.

## MATERIALS AND METHODS

### Tick collection and identification

Questing *D. reticulatus* ticks were collected in April 2021 by blanket dragging in the Pettemerduinen (52°46’46.9”N, 4°40’17.6”E), an open dune area grazed by free-ranging cattle, located in the Netherlands. Ticks were morphologically identified to species level using an identification key (Arthur, 1963; Hillyard, 1996). The collected ticks were stored at −80° C. Three of the collected *D. reticulatus* adult females were designated as biospecimen Penny1904 (Penny), Elska1004 (Elska), and Louise2209 (Louise).

### DNA extraction and library preparation

DNA extraction, library preparation, and sequencing of the three *D. reticulatus* individuals followed our previous study of *I. ricinus* Ronai et al. (2026). Briefly, 4.15-4.57 μg of DNA was extracted with the QIAamp DNA mini kit (Qiagen, Hilden, Germany). DNA quality was measured via electrophoresis using the Genomic DNA ScreenTape Analysis on an Agilent 4200 TapeStation System (Agilent Technologies Netherlands BV, Amstelveen, The Netherlands) and DNA concentration was quantified using the Qubit dsDNA HS Assay Kit on a Qubit 3.0 Fluorometer (Life Technologies Europe BV, Bleiswijk, The Netherlands). We used this DNA to prepare three 1D ligation libraries applying the Ligation Sequencing Kit SQK-LSK110 according to the manufacturer’s instructions (Oxford Nanopore Technologies, Oxford, UK).

### Sequencing and bioinformatic workflows for *de novo* whole genome assembly

Our sequencing and bioinformatic workflows are outlined in Figure 1A-1C. The three individual tick libraries were used as biological replicates to compare the efficacy between two versions of ONT flow cells, three Guppy base-caller software versions, raw read combinations, Dorado error correction, and three assembly coverage limits for Flye (Mikhail Kolmogorov, Jeffrey Yuan, Yu Lin, & Pavel A. Pevzner, 2019).

**Figure 1.**
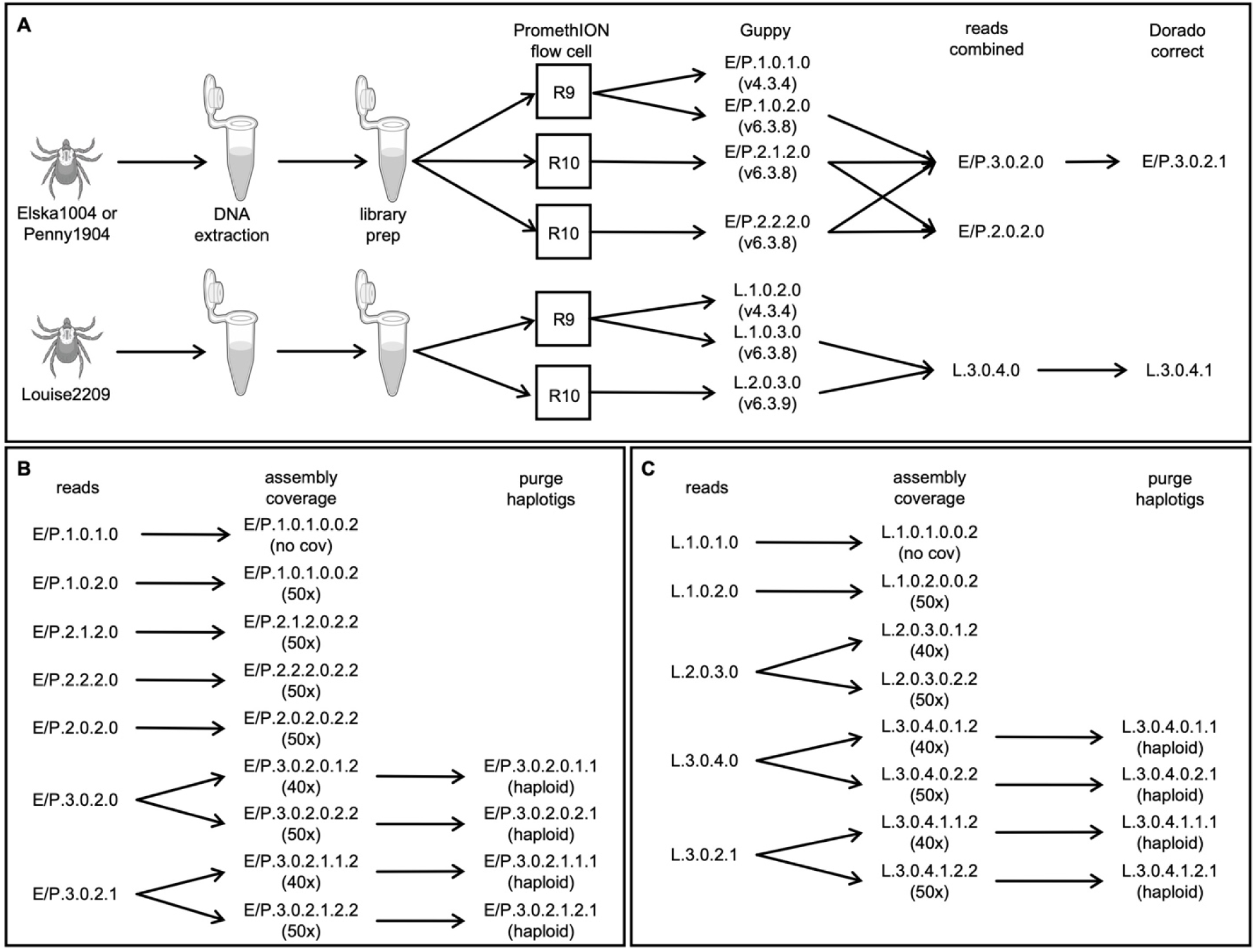
Sequencing and bioinformatic workflows for three tick samples. The sequencing and bioinformatic workflows for Elska, Louise, and Penny were similar and run in parallel. All sequence reads were assessed for quality using NanoPlot (v1.33.0, v1.42.0). All sequence assemblies were assessed for quality using QUAST (v5.2.0) and genome completeness using compleasm (v0.2.5) **A)** The sequencing and raw read workflow for Elska, Louise, and Penny consisted of six steps. After DNA extraction and library prep, the libraries were initially sequenced on a PromethION R09.4.1 flow cell with Guppy version 4.3.4 base calling, then subsequently sequenced on one or two PromethION R10.4.1 flow cells and a newer version of Guppy (v6.3.8 or v6.3.9) was used on all available data. For each individual tick, raw reads generated by the R10.4.1 flowcell and basecalled with Guppy v6.3.8 were combined. Additionally, raw reads generated by the R9.4.1 and R10.4.1 flowcells and basecalled with Guppy version 6.3.8 and 6.3.9 were combined. Raw combined reads that were basecalled using Guppy version 6.3.8 or 6.3.9 were error corrected using Dorado correct (v0.7.1). The citation for the tick icon: NIAID Visual & Medical Arts. (5/7/2025). Adult Female American Dog Tick. NIAID NIH BIOART Source. bioart.niaid.nih.gov/bioart/644. The citation for the tube icon: NIAID Visual & Medical Arts. (10/7/2024). Eppendorf Tube. NIAID NIH BIOART Source. bioart.niaid.nih.gov/bioart/143. See Table S1 for further details. **B)** The bioinformatic assembly workflow for Elska and Penny consisted of two steps. Sequence reads were assembled using Flye (v2.9, 2.9.1, 2.9.2) with no limit on assembly coverage, a limit of 40x coverage, and 50x coverage. Higher quality diploid assemblies were collapsed into their pseudo-haplotypes using purge_haplotigs (v1.1.2). See Table S2 and S4 for further details. **C)** The bioinformatic assembly workflow for Louise consisted of two steps. Sequence reads were assembled using Flye (v2.9, 2.9.1, 2.9.2) with no limit on assembly coverage, a limit of 40x coverage, and 50x coverage. Higher quality diploid assemblies were collapsed into their pseudo-haplotypes using purge_haplotigs (v1.1.2). See Table S2 and S4 for further details.

We generated 19 sets of raw reads by doing the following: Raw reads from each ONT flow cell and base-called with either Guppy version were combined, the ‘-cat’ command was used to combine raw FASTQ files. For combined raw reads that were error corrected, the Dorado correct (v0.7.1) submodule was run and output to FASTA format. All raw reads, individual and combined, were assembled using Flye (v2.9, 2.9.1, 2.9.2) (Mikhail Kolmogorov, Jeffrey Yuan, Yu Lin, & Pavel A Pevzner, 2019) using the flag ‘--genome-size 2.6g’ implemented for all assemblies and flags ‘--nano-hq’ and ‘--asm-coverage 40’ or ‘50’ for all assemblies whose raw reads were base-called using Guppy v6.3.8 and v6.3.9. Raw reads sequenced on the R09.4.1 flowcells and basecalled with Guppy v4.3.4 were excluded from raw read combining as they were considered lower quality due to the older flowcell and Guppy version.

The pseudo-haploid version of selected high quality Flye assemblies generated from combined reads was obtained using purge_haplotigs (v1.1.2) (Roach, Schmidt, & Borneman, 2018). The remaining lower quality assemblies were left diploid. First, raw reads were mapped to each diploid assembly using minimap2 (v2.26) (Li, 2018, 2021) with the flags ‘-ax map-ont’, ‘--secondary=no’ specified to create a BAM file which was sorted and indexed using SAMtools (v1.16.1) (Danecek et al., 2021) with the flags ‘-m 1G’ and -T tmp.bam’. Second, ‘purge_haplotigs hist’ was executed with the flags ‘-bam’ and ‘-genome’ to generate a coverage histogram. Third, the coverage histogram was analyzed and cutoffs identified for each diploid tick assembly by executing ‘purge_haplotigs cov’ with the flags ‘-in’, ‘-low (Elska=13, Lousie=15, Penny=5)’, ‘-mid (Elska=41, Lousie=47, Penny=36)’, and ‘-high=127’. Lastly, ‘purge_haplotigs purge’ was executed to achieve a pseudo-haploid assembly with the flags ‘-g’ and ‘-c’.

All raw reads were assessed for quality using NanoPlot (v1.33.0, v1.42.0) (De Coster & Rademakers, 2023) using the ‘-fastq’, ‘-loglength’, and ‘-N50’ flags. NanoPlot reports summary statistics specifically for raw reads. The mean and median read length and read quality is reported. The number of reads, read length N50, standard deviation read length, and total bases are reported. The number of bases at Phred quality score cutoffs greater than Q5, Q7, Q10, Q12, Q15 are reported for raw sequence reads prior to Dorado correct. Quality control statistics for all 38 diploid and pseudo-haploid Flye assemblies were generated by executing QUAST-LG (v5.2.0) (Mikheenko, Prjibelski, Saveliev, Antipov, & Gurevich, 2018) with the flag ‘-large’ specified (Mikheenko et al., 2018). For each assembly, the number of contigs, total length, largest contig length, total length, GC%, N50, N90, auN, L50, L90, and the number of Ns per 100 Kbp is reported. Raw read coverage was calculated for all raw reads before and after Dorado raw read error correction. Assembly completeness was assessed for each diploid and pseudo-haploid assembly by using compleasm (v0.2.5) (Huang & Li, 2023) under the ‘run’ submodule with the arthropoda_odb10 lineage specified. A total of 1,013 BUSCO genes from the phylum Arthropoda (arthropoda_odb10) were used to assess each assembly. The number and percentage of complete, single, duplicate, fragmented, and missing BUSCOs are reported.

### Contaminant identification and removal

Contaminants within the three chosen pseudo-haploid assemblies to represent each tick were identified and removed using the NCBI Foreign Contamination Screen (v0.5.0) (Astashyn et al., 2024) adaptor (FCS-adaptor) and genome cross-species (FCS-GX) tools. For FCS-adaptor, the ‘-euk’ tag was used and for FCS-GX, tax-id 34619 was used as there were no *D. reticulatus* assemblies publicly available at the time of screening. Four additional contaminants were found and removed from the soft-masked Penny assembly during submission to NCBI GenBank. Once contaminants were removed, assembly quality metrics using QUAST-LG (v.5.2.0) and completeness metrics using compleasm (v.0.2.8) were assessed for each assembly.

### BRAKER3 gene prediction

We annotated the protein-coding genes in the *D. reticulatus* Louise pseudo-haploid assembly using BRAKER3 (v3.0.8) (Brůna, Hoff, Lomsadze, Stanke, & Borodovsky, 2021; Brůna, Lomsadze, & Borodovsky, 2024; Buchfink, Xie, & Huson, 2015; Gabriel et al., 2024; Gabriel, Hoff, Brůna, Borodovsky, & Stanke, 2021; Gotoh, 2008; Hoff, Lange, Lomsadze, Borodovsky, & Stanke, 2016; Hoff, Lomsadze, Borodovsky, & Stanke, 2019; Iwata & Gotoh, 2012; Kim, Paggi, Park, Bennett, & Salzberg, 2019; Kovaka et al., 2019; Pertea & Pertea, 2020; Stanke, Diekhans, Baertsch, & Haussler, 2008; Stanke, Schöffmann, Morgenstern, & Waack, 2006). Then we downloaded raw read RNA-seq data from NCBI: *D. reticulatus* salivary glands from whole larvae (SRR950367) (Villar et al., 2014), adult males (SRR9640925 and SRR9640926) (Kartashov et al., 2023), and adult females (SRR9640927 and SRR9640928) (Kartashov et al., 2023) using the prefetch command in SRA Toolkit (v3.2.0) (https://github.com/ncbi/sra-tools). Protein data was downloaded from the OrthoDB (v.11) (Kuznetsov et al., 2023) Arthropoda library.

The AUGUSTUS gene annotation was chosen over the BRAKER3 annotation, as the gene structure most closely resembled that of other *Dermacentor* assemblies (Billows et al., 2026; Cassens et al., 2025; Jia et al., 2020; Tompkin et al., 2026). The AUGUSTUS GTF file was transformed to GFF3 using the gtf2gff.pl Perl script (v3.2.1) (https://github.com/nextgenusfs/augustus) provided by the creators of AUGUSTUS (with the flags --gff3, --printExon, --printUTR, --printIntron) to generate a file that follows the parent-child hierarchy. The subsequent GFF3 file representing the Louise annotations was used in Liftoff (v1.6.3) (Shumate & Salzberg, 2021) to identify genetic features in *D. reticulatus* Elska and Penny pseudo-haploid assemblies. Genetic features were counted in each pseudo-haploid assembly GFF3 file using the following on command line: cut -f 3 file.gff3 | sort | uniq -c. The completeness of the Louise protein FASTA was assessed by executing compleasm (v0.2.6) (Huang & Li, 2023) under the ‘protein’ submodule with the arthropoda_odb10 lineage specified.

### Ortholog gene synteny of whole genome assemblies from *Dermacentor* spp

We identified and visualized orthologous, syntenic regions between the transcripts and longest isoform per gene for six *Dermacentor* assemblies.

For the transcript syntenic analysis we performed the following steps. We obtained the genomic GFF and translated CDS files from the following four NCBI RefSeq IDs: GCF_013339745.2 (*D. silvarum*) (Jia et al., 2020), GCF_038994185.2 (*D. albipictus*), GCF_023375885.2 (*D. andersoni*) (Tompkin et al., 2026), and GCF_050947875.1 (*D. variabilis*) (Cassens et al., 2025). We note that the *D. albipictus* assembly (GCA_038994185.2) represents a haplotype from a single male tick, *D. silvarum* (GCA_013339745.2) was assembled from a mixture of tick larvae (siblings), and the remaining assemblies are each from a single female. The NCBI files were parsed into their respective BED and peptide directories which was performed by executing GENESPACE (v1.2.3) (Lovell et al., 2022). The pseudo-haploid Louise assembly was scaffolded with RagTag (v2.1.0) (Alonge et al., 2022) to improve contiguity and syntenic comparisons between genomes, and re-annotated with BRAKER3, as previously described. The ‘scaffold’ command was used to execute RagTag where the chromosome-sized *D. reticulatus* assembly (Devon; Billows et al., 2026) was used as the reference. The AUGUSTUS GTF for Louise and the GALBA GTF for Devon were filtered to include only transcripts and then converted to the GENESPACE BED file format and moved to the same BED directory mentioned in the first step. Both amino acid FASTA files for Louise and Devon were moved to the peptide directory mentioned in the first step.

For the longest isoform syntenic analysis we performed the following steps. The longest isoform per protein-coding gene per *Dermacentor* genome was extracted, with the goal of reducing synteny of repeated transcripts. Genome annotations for the six previously mentioned *Dermacentor* tick assemblies were preprocessed using AGAT (v1.4.2) (Dainat, 2022) using the ‘agat_sp_keep_longest_isoform.pl’ command and ‘-gff’ flag to extract the longest isoform per gene. Protein sequences were extracted from each filtered annotation using gffread (v0.12.7) (Pertea & Pertea, 2020) with ‘-S’ and ‘-y’ flags to translate coding sequences.

For both syntenic analyses we performed the following steps. GENESPACE-formatted BED files were generated from the processed GTF files by parsing transcript features. Gene ID concordance between BED and peptide FASTA files was verified for all six assemblies prior to downstream analysis. Orthogroups were inferred from protein sequences using OrthoFinder (v2.5.5) (Emms & Kelly, 2019) and supplied as input to GENESPACE (v1.2.3) for syntenic block detection and riparian plot generation, using the *D. reticulatus* Devon assembly as the reference genome given its Hi-C level scaffolding of chromosomes. GENESPACE accessed DIAMOND2 (v2.18) (Buchfink, Reuter, & Drost, 2021) and MCScanX (v1.0.0) (Yupeng Wang et al., 2012). Each species assembly was plotted in order according to the OrthoFinder species tree.

Finally, inverted chromosome-sized scaffolds relative to the reference were identified by visual inspection of braid crossing patterns in initial plots and corrected iteratively using the ‘invertTheseChrs’ parameter. Reference-guided chromosome assignments consistent with synteny were manually re-ordered and re-oriented to maximize alignment. The width of each syntenic block, or chromosome-sized scaffold, represents genes that stay in the same order across two genomes and are numbered #1-11, except for *D. albipictus* which was numbered #1-10. The synteny plots were generated using scripts available here: https://github.com/katiecdillon/Dreticulatus/tree/main/synteny_plots.

### Shared orthologs between three *Dermacentor* assemblies

We identified unique and shared orthologous genes between two closely related *Dermacentor* species using three assemblies: *D. reticulatus* Devon, *D. reticulatus* Louise, and *D. silvarum* (Jia et al., 2020). We include Devon in this analysis to visualize similarities and differences between the two *D. reticulatus* assemblies.

Orthologous gene groups were identified using OrthoFinder (v3.1.0) and protein sets from the three tick assemblies in GFF3 format were included in the analysis. GFF files were first cleaned with an in-house Python (v3.x Google compute engine backend) script to remove any unknown, invalid or non-permissible characters. FASTA IDs were changed to clean headers to account for errors in differences of annotation methods. OrthoFinder performed an all-versus-all protein sequence similarity search using DIAMOND to identify homologous proteins among the proteomes. It then created a similarity graph which was clustered into orthogroups using a Markov Clustering algorithm, and gene trees were subsequently inferred to differentiate orthologous and paralogous relationships. Pairwise shared orthogroups were also identified.

Counts of shared and unique orthogroups were summarized and visualized as Venn diagrams. The three-way Venn diagram was visualized using the Python package matplotlib-venn, pandas, and numoy via an in-house Python (v3.x Google compute engine backend) script. The custom script imported the OrthoFinder orthogroups.tsv file into a pandas DataFrame. Each orthogroup was evaluated for the presence or absence of proteins within the three proteomes. Based on this pattern, orthogroups were assigned to the categories: unique, pairwise shared, or universally shared orthogroups. The script then summarized the number of orthogroups in each category and exported category-specific orthogroup lists for diagram creation. The scripts used to generate the values and plot for the Venn diagram can be accessed here: https://github.com/katiecdillon/Dreticulatus/tree/main/venn_plot.

### Manual transposable element characterization

Manual curation of TEs in tick genome assemblies is the gold standard, as automated curation misses a large proportion of TE diversity (Dillon et al., 2026). However, manual curation of TEs is complex, as previously identified consensus sequences can be mistakenly re-identified, wasting time and effort. Therefore, we developed a novel pipeline to manually annotate the TEs in our reference *D*. *reticulatus* pseudo-haploid assembly Elska. The Elska assembly was chosen due to its low duplicate BUSCO score.

The tool HiTE (Hu et al., 2024) was used to generate a preliminary library for Elska. Then we implemented TECurate (https://github.com/davidaray/bioinfo_tools/blob/master/template_TEcurate.sh), a custom pipeline that uses TEAid (Goubert et al., 2022) and custom scripts (https://github.com/katiecdillon/Dreticulatus/tree/main/TE_curation) to generate a preliminary identification of a consensus sequence’s TE class (LINE, LTR, DNA transposon, Rolling Circle, etc.) and TE superfamily (LINE1, Gypsy, etc.), using the presence of known TE-derived open reading frames. Potential TE consensus sequences without open reading frames (SINEs, non-autonomous DNA transposons, etc.) were generated by HiTE, requiring additional visual examination of TEAid output plots of divergence, coverage, blastn dotplots, and structure/protein hits. In this step, output files associated with each consensus sequence was examined for hallmarks of known TE types (see examples at the TE-Aid github site) and assigned a likely TE Class and superfamily, if possible. If no clear hallmarks were identifiable, they remained in the Unknown category. A TE library in FASTA format was formed from the consensus sequences assigned with a likely TE class and Unknown TEs.

To confirm TE class, family, and identity, the TE library was submitted to the TEClass2 (Bickmann, Rodriguez, Jiang, & Makalowski, 2026) web portal. The database of identities with probabilities calculated at over 70% for each TE consensus sequence was then used to alter headers in the existing library. This library was then used as the query for RepeatMasker runs on each of our three *D. reticulatus* assemblies (Elska, Louise, and Penny) and this output was used as input for the scripts we used to generate plotted data. Plots were generated using ggplot2. Total TE-derived genome volume (bp) was calculated using RepeatMasker output for each range of divergences. Afterward, the proportion derived from each Class and, within each Class, each major Superfamily was calculated to allow comparison of relative rates of accumulation in each divergence range. The highest quality assembly for each tick was identified, and the assembly removed of contaminants and soft-masked for repeats using the manually curated repeat library.

### Variation within and among individuals

We compared inter- and intra-genome nucleotide diversity of the three *D. reticulatus* samples from two populations: The “Netherlands” (n=3; Elska, Louise, Penny) and the “UK” (n=1); all pseudo-haploid assemblies. Two assemblies were used as references: The Devon assembly acquired from NCBI (GCA_057332255.1) and the Louise assembly (L.3.0.4.1.1.1), of which repeats were soft-masked and contaminants removed as described previously. Devon represented a single sample within the “UK” population and Elska, Louise, and Penny represented three samples within the “Netherlands” population. Raw read IDs E.3.0.2.0, L.3.0.4.0, P.3.0.2.0, and combined Devon raw reads (50 files; SRR38614766–SRR38614815) were error corrected using Dorado correct (v2.0.0). Dorado correct output is only available in FASTA format. Next, corrected raw reads were aligned to both reference assemblies using minimap2 (v2.29) and output to a SAM file. SAMtools (v1.21) was used to convert to BAM format, sort the BAM files, and index them.

For the raw read files aligned to the Devon assembly, SNPs and indels were identified using BCFtools mpileup (v1.23.1) to summarize base calls for each of the 11 largest (chromosome-sized) scaffolds of Devon, variants called, and output to a VCF file. For the raw read files aligned to the Louise assembly, minimap2 was used to align the Louise assembly to the Devon assembly and SAMtools used to convert to BAM format, sort the BAM file, and indexed. Using SAMtools, the alignment BAM file was used to extract the Louise assembly contigs that mapped to the 11 chromosome-size scaffolds of Devon and output to a regions file containing Louise contigs that mapped to each chromosome-sized scaffold. Variants were then identified using BCFtools mpileup to summarize base calls for each of the regions files, variants called, and output to a VCF file.

All VCF files underwent site-level filtration using BCFtools where indels, all missing genotypes, and sites with total depth outside of 20x-300x were excluded, and CSI indexed. Filtration for total depth was determined by plotting the INFO/DP column of each VCF file for upper and lower quantiles (Figure S1). For INFO/DP < 20 we aimed to exclude low-coverage sites and for INFO/DP >300 we aimed to exclude repetitive loci (Figure S1). The population genetics statistic summary tool, Pixy (v2.2.0) (Bailey, Stevison, & Samuk, 2025; Korunes & Samuk, 2021), was used on all 22 filtered VCF files. The summary statistics pi, Watterson’s theta, Tajima’s D, Hudson’s F_ST_, and d_xy_ were calculated for both populations and each chromosome-sized scaffold. Pi and Watterson’s theta summary statistics were calculated for each chromosome-sized scaffold and split between the two populations. The d_xy_ and Hudson’s F_ST_ summary statistics were calculated for each chromosome-sized scaffold for both populations combined. The Tajima’s D summary statistic was calculated for each chromosome-sized scaffold for the Netherlands population only. The window size was set at 50 Kbp for each population, based on the low sample size and need for increased resolution (Subramanian, 2016). We followed the post-hoc aggregation steps outlined in the Pixy documentation for all summary statistics genome wide and for each chromosome-sized scaffold (Bailey et al., 2025).

The number of individuals used here is far too low for in depth population genomic analyses and is well below the sampling used in population genomics of other ticks (Dong & Schoville, preprint; Frederick et al., 2023; Schoville et al., 2024). However, we are interested in an initial estimate of genetic diversity, and some measures, such as pi, are not biased due to sample size [vs. measures such as theta or calculations dependent upon theta, such as Tajima’s D (Subramanian, 2016)]. The scripts used to generate this data are located here: https://github.com/katiecdillon/Dreticulatus/tree/main/pixy_plots.

## RESULTS

### Raw read quality improves after Dorado error correction

For the three *D. reticulatus* ticks we sequenced, the 19 sets of raw reads (Figure 1A) varied in amount and quality (Table S1). The average number of non-combined, uncorrected raw reads was ∼14.3 million, compared to ∼33 million combined, uncorrected raw reads (Table S1). The average number of raw reads basecalled by Guppy v4.3.4 was ∼13.7 million and ∼16.6 million by Guppy v6.3.8 and v6.3.9 both (Table S1). Prior to Dorado error correction, the total average number of reads from combined read sets was ∼33 million and after error correction, ∼7.4 million reads; ∼ 78% decrease in total reads (Table S1). In contrast, following Dorado error correction, an average of 74.4% (range 72.2% - 76.4%) of bases were retained, with an increase of mean read length of ∼12.2 Kbp (range 11.5 – 12.8 Kbp) of reads after error correction (Table S2). Thus, Dorado error correction removed a large number of the shortest reads.

### Assembly parameter comparisons reveal high-quality assemblies

We assessed the 38 diploid and pseudo-haploid assemblies generated from the three *D. reticulatus* ticks. Assemblies generated from combined raw reads had ∼52% fewer contigs, ∼44% larger contig N50, and ∼1% more complete BUSCOs compared to reads from individual flow cells (Figure 2A-D, Table S3-S4). We then made four comparisons to determine the impact of major parameters on the resulting assemblies.

**Figure 2.**
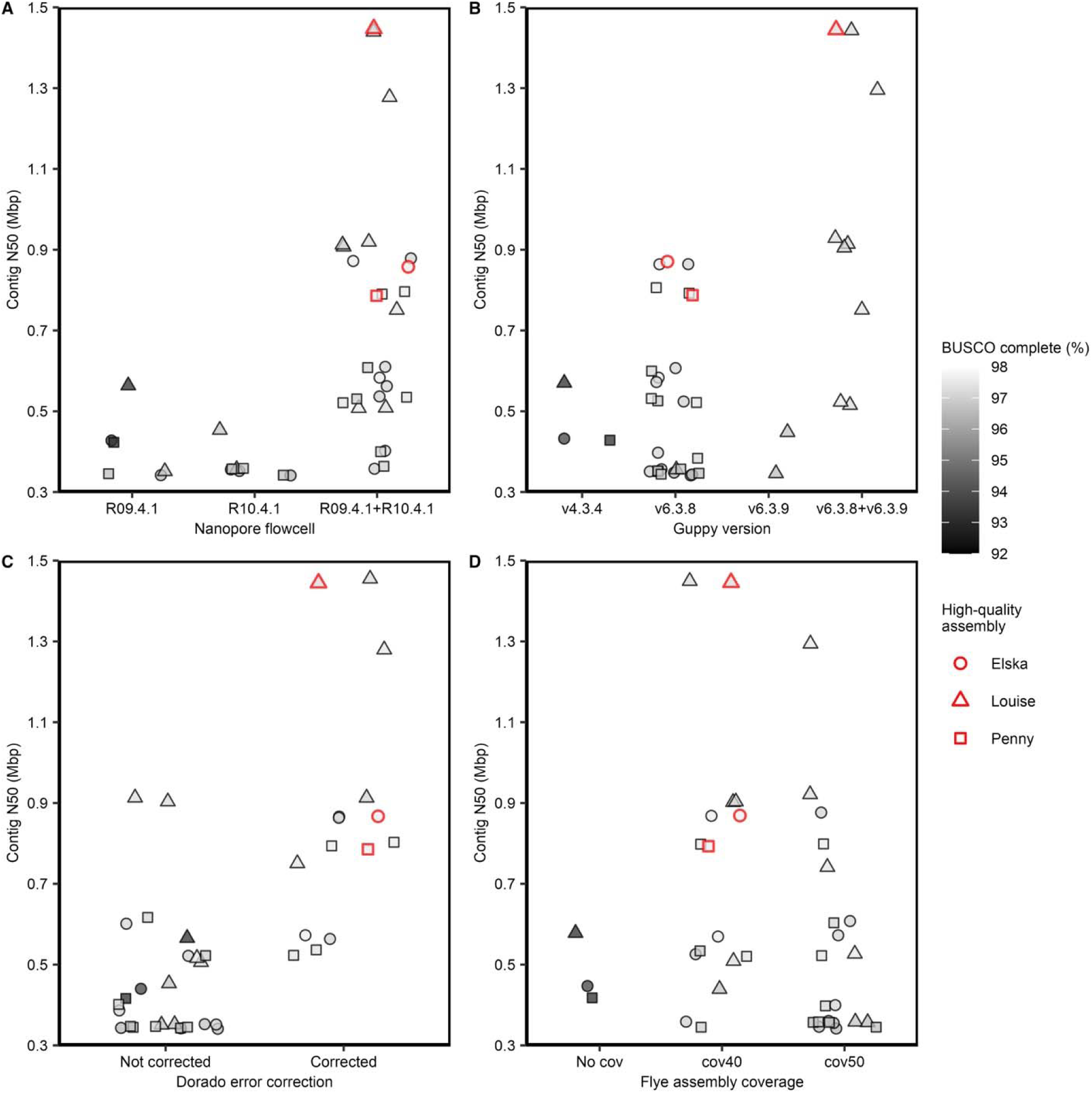
Quality assessment of 38 genome assemblies across test variables. Each point is a genome assembly, with the highest quality assembly for each tick individual denoted in red. Quality of the assembly was assessed with contig N50 Mbp (the length of the shortest contig that must be included to cover half an assembly), as N50 increases, fewer longer contigs are used to cover half of the assembly, leading to a higher-quality assembly. BUSCO complete score (the sum of the single copy and duplicated genes that can be completely aligned to the assembly with one or more than one gene copy present), a higher BUSCO score indicates higher assembly completeness and a higher quality assembly. **A)** Nanopore flow cell, a device where long-read sequencing occurs, R09.4.1, R10.4.1, and combined R09.4.1+R10.4.1. **B)** Guppy base caller, a data processing tool that can convert raw signal data generated by the flow cell into DNA sequence data, v4.3.4, v6.3.8, v6.3.9, and combined v6.3.8+v6.3.9. **C)** Raw reads, the DNA sequence data, not corrected and raw reads corrected with Dorado. **D)** Flye, a *de novo* bioinformatic assembler designed for long-read single molecule sequencing reads, assembly coverage to target coverage for the longest reads, with no limit on assembly coverage, a limit of 40x coverage, and 50x coverage. See Table S2 and S4 for further details.

First, we assessed the assemblies from two different types of flow cells (R09.4.1 and R10.4.1). Between individual flow cells, the average number of contigs increased from R09.4.1 to R10.4.1 by ∼16%, contig N50 decreased by ∼12%, and complete BUSCO increased by ∼1.8% (Figure 2A, S3-S4). Between individual and combined flow cells, the number of contigs decreased from individual to combined by ∼48%, contig N50 increased by ∼56%, and complete BUSCO increased by ∼1.01% (Figure 2A, Table S3-S4).

Second, we assessed the assemblies whose reads were base called by three different versions of Guppy (Figure 2B, Table S3-S4). Between read sets that were base-called by Guppy v4.3.4, v6.3.8, and v6.3.9 the average number of contigs decreased between Guppy v4.3.4 and v6.3.8 by ∼14% and increased between Guppy v6.3.8 and 6.3.9 by ∼46%, contig N50 increased by ∼4% between Guppy v4.3.4 and v6.3.8 and decreased between 6.3.8 and 6.3.9 by ∼20%, and complete BUSCO increased between Guppy v4.3.4 and v6.3.8 by ∼4% and between v6.3.8 and v6.3.9 decreased by ∼1% (Figure 2B, Table S3-S4). Between read sets base called by individual Guppy versions (v.6.3.8 and v6.3.9) and combined Guppy versions (v6.3.8+v6.3.9), the number of contigs between individual and combined Guppy versions decreased by ∼36%, contig N50 increased by ∼46%, and complete BUSCO increased by ∼1% (Figure 2B, Table S3-S4).

Third, we assessed the assemblies whose reads were either not corrected or were error-corrected by Dorado (Figure 2C, Table S3-S4). Assemblies with error-corrected reads had on average ∼51% fewer contigs, ∼44% larger contig N50, and ∼1% increase in complete BUSCO (Figure 2C, Table S3-S4).

Fourth, we assessed the assemblies where three Flye assembly coverages were applied (Figure 2D, Table S3-S4). Assemblies generated from all reads, regardless of coverage, had ∼31% more contigs, ∼29% smaller contig N50, and ∼4.1% fewer complete BUSCO than assemblies with a maximum of 40x assembly coverage. Assemblies with a maximum of 40x assembly coverage had ∼25% fewer contigs, ∼21% larger contig N50, and ∼0.3% larger complete BUSCO than assemblies with 50X maximum assembly coverage (Figure 2D, Table S3-S4). The difference between 40x and 50x maximum coverage is 40x maximum favors longer reads.

In summary, the highest quality pseudo-haploid assemblies generated for the three individual ticks were obtained from raw combined reads sequenced on an R09.4.1 and R10.4.1 flow cell, base-called by Guppy v6.3.8 and v6.3.9, error corrected with Dorado and assembled with Flye using 40x maximum coverage. One pseudo-haploid assembly was selected to represent each tick based on quality and assembly completeness metrics. For the tick Elska, the assembly E.3.0.2.1.1.1 was selected, for Louise the assembly was L.3.0.4.1.1.1, and for Penny assembly P.3.0.2.1.1.1 was selected (Figure 2A-D, Table S3-S4). On average, the length of the three assemblies is ∼2.29 Gbp. The list of bacterial and adaptor contaminants is reported in Table S5 and S6. The final, pseudo-haploid assemblies with contaminants removed have on average 7,353 contigs, 1.03 Mbp N50, 95.92% complete BUSCO, and 0.84% duplicate BUSCO (Table S7).

### Gene annotation of the Elska, Louise, and Penny assemblies

We selected the AUGUSTUS gene predictions over the BRAKER3 gene predictions as they more closely resembled total gene counts from other ticks (Billows et al., 2026; Cassens et al., 2025; Jia et al., 2020; Tompkin et al., 2026). On average, we detected ∼41,953 protein-coding genes for the three assemblies. For the pseudo-haploid Louise assembly, the protein BUSCO score was ∼95.92% (79.78% single, 16.14% duplicate). Among the three new *D. reticulatus* assemblies, the number of genes was similar to those reported for other *Dermacentor* species genomes (Billows et al., 2026). The complete protein BUSCO score was similar to the *D. reticulatus* Devon assembly, but the duplicate protein BUSCOs in Louise are much higher, indicating that additional work is needed on the annotation (File S1). As the Louise genome is fragmented, there may be missing orthologs.

### Orthologous gene synteny assigns scaffolds to *D. reticulatus* chromosomes

We compared gene synteny across all transcripts (Figure 3) and across the longest isoform per gene (Figure S2) for the six available *Dermacentor* assemblies. For all transcripts among the six assemblies, there is a total of 48,604 orthogroups, of which 10,586 orthogroups containing 101,474 transcripts are shared across all six *Dermacentor* assemblies (Figure 3). For the longest isoform per gene across all six assemblies, there were 166,495 genes total, with a mean protein length of ∼425 amino acids (median 314 amino acids) (Figure S2). There is a total of 39,461 orthogroups, of which 10,970 orthogroups containing 71,983 genes are shared across all six assemblies (Figure S2). These ribbon plots clearly indicate which chromosome-sized scaffolds contain the most shared orthogroups. In Figure 3 and Figure S2, chromosome 1 for *D. albipictus* is a fusion of chromosomes 1 and 2 of all other *Dermacentor* species. The ribbon plots also facilitate a clear understanding of orthogroups among the assemblies and species, strongly suggesting chromosome assignments of scaffolds among these species. For both the gene synteny across transcripts (Figure 3) and across the longest isoforms per gene (Figure S2), the chromosome width is scaled by gene rank order (number of genes) and both *D. reticulatus* assemblies have roughly twice as many genes.

**Figure 3.**
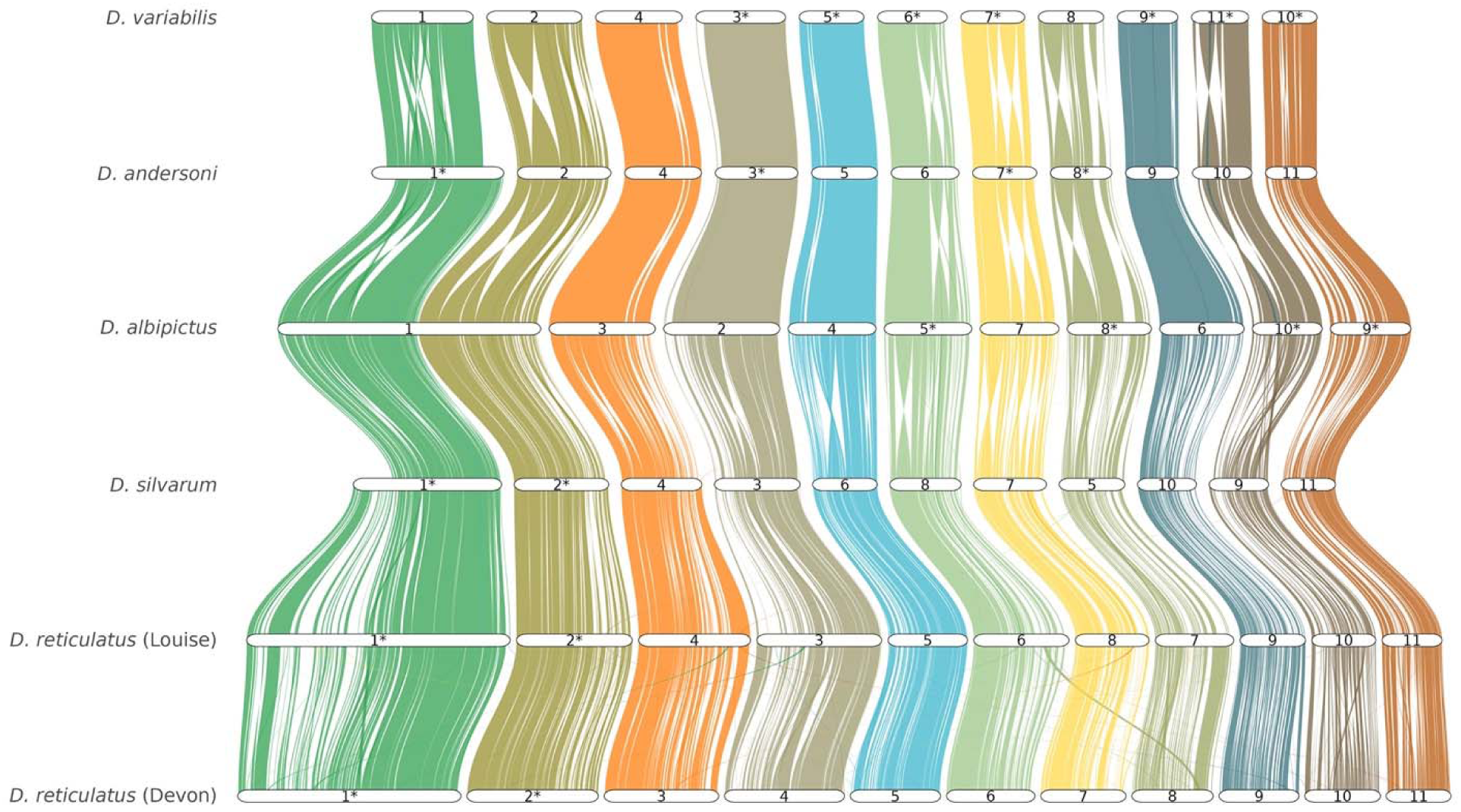
Gene synteny of transcripts in six *Dermacentor* assemblies. The six tick assemblies are ordered according to the OrthoFinder species tree starting with the 11 chromosome-sized scaffolds of *D. reticulatus* (Devon). Chromosome-sized scaffolds are numbered according to size from left to right and the transcripts shared between each scaffold given a unique color. Chromosome-sized scaffolds were ordered to minimize crossing of ortholog sets and inverted chromosomes (note: *D. albipictus* scaffold 1 is shown crossing to minimize). Asterisks (*) denote the scaffolds that were manually flipped in GeneSpace.

The closest relative to *D. reticulatus* is *D. silvarum* (Billows et al., 2026), so we compared the orthogroups among *D. reticulatus* (Louise), *D. reticulatus* (Devon), and *D. silvarum* assemblies (Figure 4). A total of 15,766 orthogroups are shared among all three *Dermacentor* assemblies, with both *D. reticulatus* assemblies sharing 5,750 orthogroups not found in *D. silvarum* (Figure 4). Each *D. reticulatus* assembly shares between 920 to 1,487 orthogroups with *D. silvarum*, however each *D. reticulatus* assembly has more than 600 unique orthogroups (Figure 4). All specified unique and shared orthogroups are in File S2.

**Figure 4.**
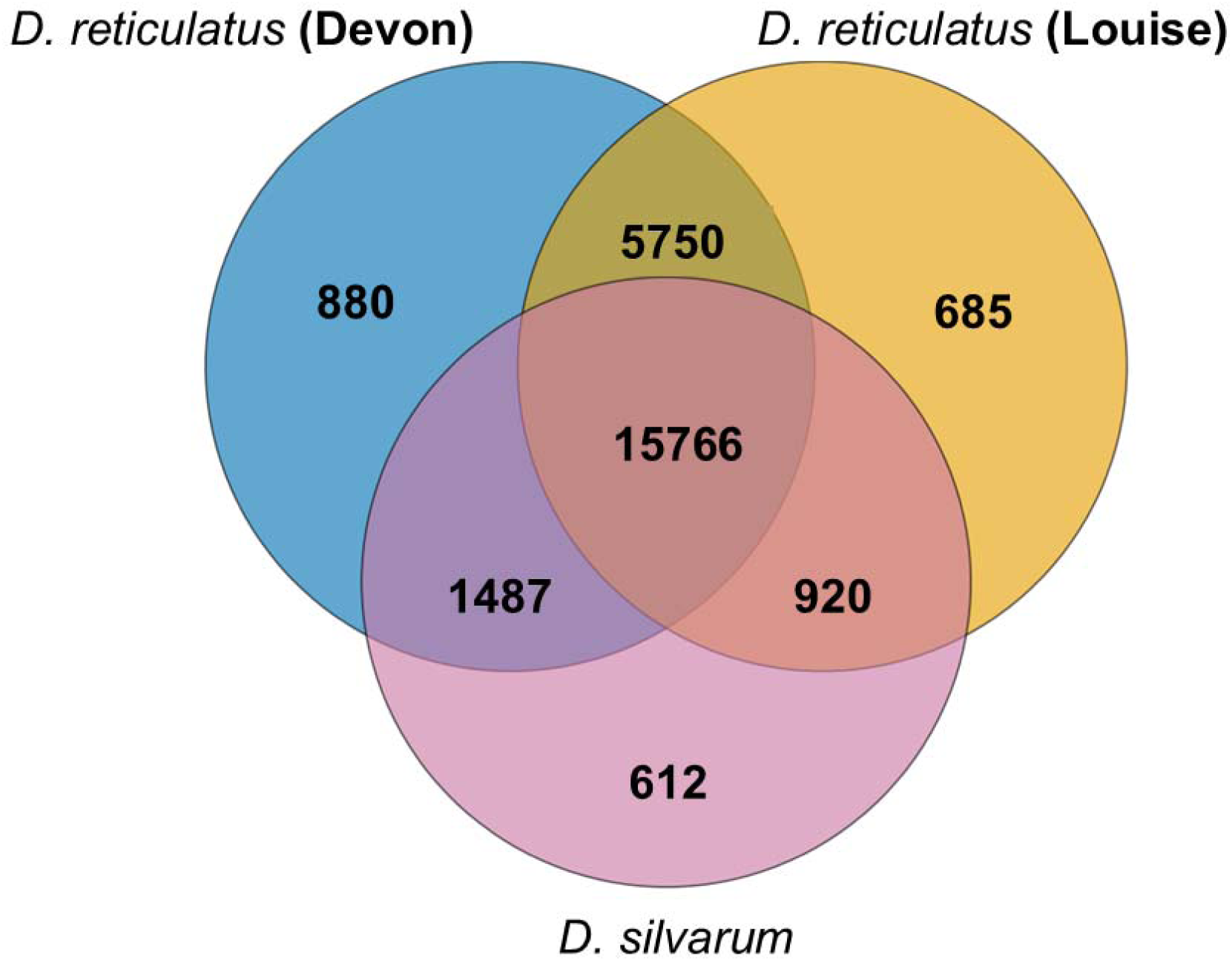
Orthogroups shared among three *Dermacentor* genome assemblies. The number of orthologous gene groups identified by OrthoFinder that are unique to, or shared among, *D. reticulatus* (Devon), *D. reticulatus* (Louise), and *D. silvarum* (BIME_Dsil_1.4). Numbers in each section indicate the count of orthogroups present in the corresponding genome assembly or assemblies. The central intersection represents orthogroups universally shared across all three assemblies.

### Transposable elements in *D. reticulatus*

The pseudo-haploid Louise assembly contains 63.3% TEs (Figure 5A, Table S8), Elska contains 63.2%, and Penny contains 63.5% (Figure S3A, Table S8). The TEs that dominate the three assemblies are LTR elements ranging from 20.98% to 21.21%, DNA from 14.86% to 14.96%, and LINE 9.03% to 9.06% (Figure 5A, Figure S3A, Table S8). Unknown TEs range from 16.47% to 16.54% (Figure 5A, Figure S3A, Table S8). From these data we see that between species, on average, *D. reticulatus* Elska, Louise, and Penny have 7.07% and 8.03% fewer total TEs, 59.08% and 59.46% fewer DNA transposons, 40.45% and 39.74% fewer LINE elements, and 50.71% and 59.01% more LTR elements than *I. ricinus* and *I. scapularis*, respectively (Ronai et al., 2026).

**Figure 5.**
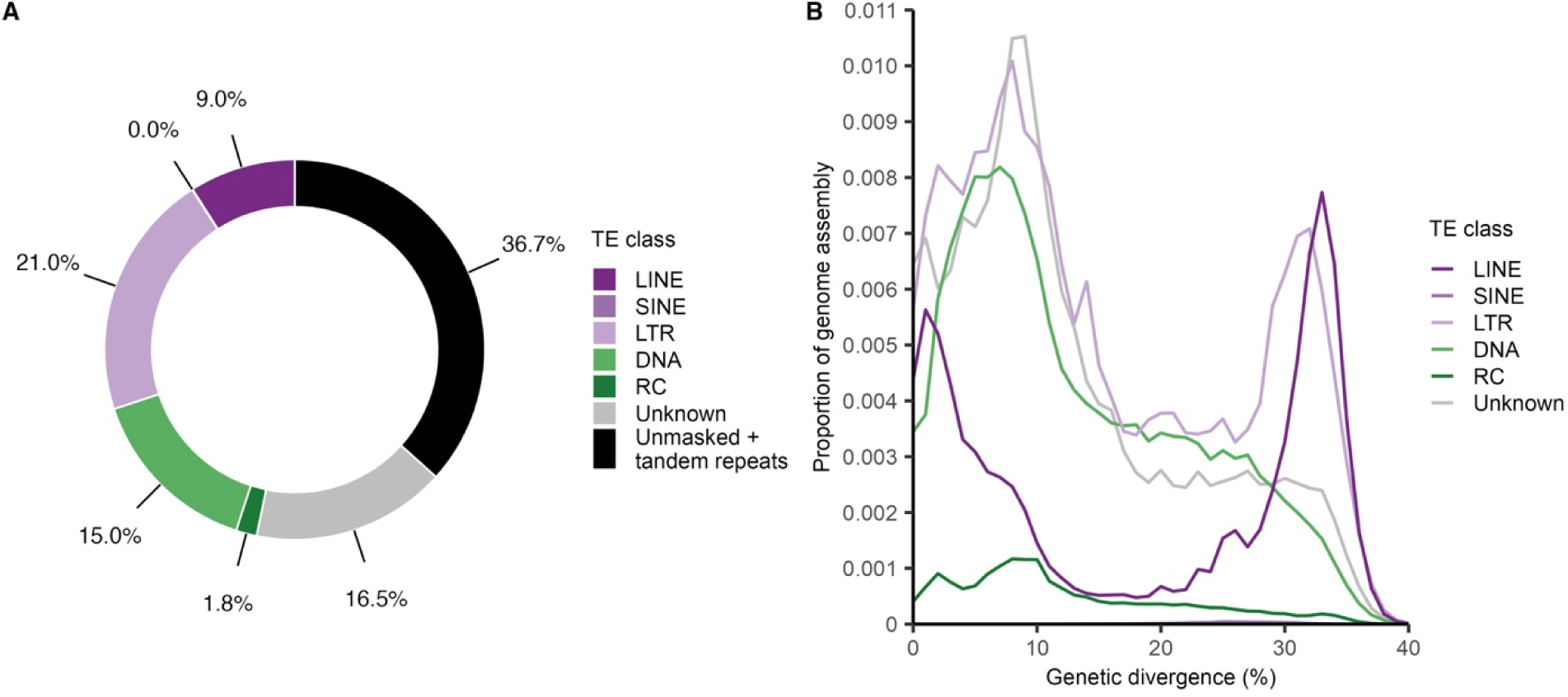
Characterization of the transposable elements in *D*. *reticulatus* Louise. The transposable element landscape of *D. reticulatus* Louise. **A)** Proportion of the Louise assembly occupied by the major TE classes, unknown TEs, tandem repeats, and unmasked sequences. **B)** Repetitive proportion of TE classes and genetic divergence in the Louise assembly.

LINE and LTR elements peaked at two distinguishable divergences in the three assemblies, LINE and LTR retrotransposons share a of peak of amplification at the higher divergence bins (∼32-33%), while in the more recent past (0-12%), LINEs appear to be on the increase and LTR retrotransposons have waned in their accumulation (Figure 5B, Figure S3B). DNA transposons with Terminal Inverted Repeats (TIRs) such as hAT and TcMariner transposons experienced a gradual increase until recently and, like LTR retrotransposons are waning in their accumulation but are still impacting genome structure (Figure 5B, Figure S3B). Rolling Circle transposons appear to have maintained a low level of activity over the evolution of the *D. reticulatus* genome and SINEs play an essentially non-existent role (Figure 5A-B, Figure S3A-B).

We also compared changes in TE accumulation overall and across time in both clades. When comparing overall genome content, *D. reticulatus* individuals exhibit substantially lower DNA transposon accumulation when compared to their Ixodid counterparts (Figure S4A-B). This is reflected in the heatmap (Figure S4C) via a large increase in DNA transposon accumulation in the recent past in *Ixodes* ticks while LINEs have flourished more in *Dermacentor*. SINEs appear to have decreased substantially in both species and in both lineages (Figure S4A-C). This is a proportional decrease and, in the context of very low overall SINE content (Figure 5A-B, Figure S3A-B, Table S8), is an artifact of this general lack of SINE activity.

The final, pseudo-haploid assemblies with contaminants removed and soft-masked with the manually curated TE library were uploaded to NCBI GenBank. We observed relatively few differences in the amount or composition of TE classes among individual *D. reticulatus* assemblies. There is a maximum of 0.3% difference in TEs between all three *D. reticulatus* assemblies (Figure 5A, Figure S3A).

### Variation within and among individuals of *D. reticulatus*

We compared inter- and intra-genome nucleotide diversity of the three *D. reticulatus* samples from two populations: The “Netherlands” (n=3; Elska, Louise, Penny) and the “UK” (n=1). When the Louise assembly was used as the reference, the average pi for the Netherlands population was 0.0149 and the UK was 0.0132 (Figure 6A-B). Watterson’s theta, Tajima’s D, d_xy_, and Hudson’s F_ST_ statistics were also calculated with Louise as the reference (Figure S5A-C). The average d_xy_ between the Netherlands and UK populations was 0.0177 (Figure S5C), and the average F_ST_ (variance between populations) was 0.266 (Figure S5D). When the Devon assembly was the reference, the average pi for Netherlands was 0.0155 and the UK was 0.0102 (Figure 6C-D). Watterson’s theta, Tajima’s D, d_xy_, and Hudson’s F_ST_ statistics were also calculated with Devon as the reference (Figure S6A-C). The average d_xy_ for the Netherlands and UK populations was 0.0173 (Figure S6C) and the average F_ST_ was 0.302 (Figure S6D). Summary statistics were also calculated per chromosome for each reference assembly (Table S9A-S9C).

**Figure 6.**
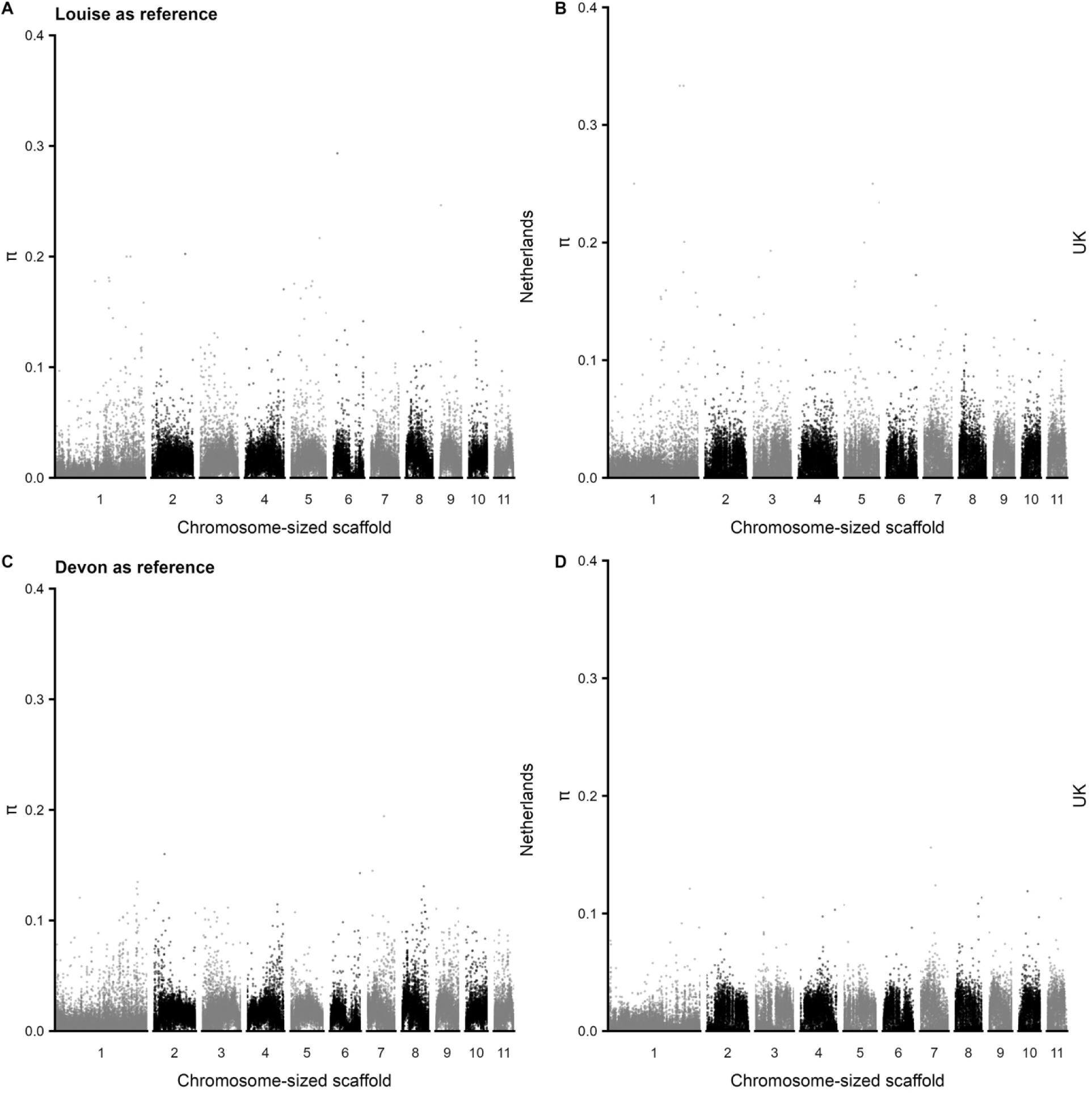
Genome-wide population pi values for *D*. *reticulatus* Netherlands and UK ticks. The “Netherlands” population consists of three tick biosamples: Elska, Louise, and Penny (panels A, C). The “UK” population consists of one tick biosample: Devon (panels B, D). **A, B)** The Louise contigs were mapped to the eleven chromosome-sized scaffolds of the Devon assembly and were then used as the reference for alignment and SNP calling. **C, D)** The Devon assembly was used as the reference.

## DISCUSSION

We establish best practices to generate high-quality genome assemblies for large, highly repetitive animal genomes using a widely used sequencing platform for molecular ecology studies. Our assessment of the 38 diploid and pseudo-haploid genome assemblies we generated established an optimal path for genome assembly of deeply sequenced tick genomes using ONT, novel characterization of TEs in *D. reticulatus* ticks, and an estimated number of chromosomes for *D. reticulatus*.

We examined the effects of ONT sequencing flow cell versions, Guppy base-calling versions, and raw read combinations to inform the best path forward for tick genome assembly. The data were collected before and after ONT made substantial advances in their nanopores and software, going from nanopores with single read points (R09.4.1) and limited base-calling software (Guppy v4.3.4) to longer nanopores with two read points (R10.4.1) and improved base-calling (Guppy v6.3.x). As expected, we find combining raw reads base-called by later Guppy versions results in a higher raw read yield than using single-run raw reads. In particular, our use of the Dorado raw read error correction subcommand greatly improves raw read quality by discarding short and low quality reads and re-implementing the HERRO algorithm, the deep learning model designed to improve nanopore read accuracy (Stanojević, Lin, Nurk, Florez de Sessions, & Šikić, 2026).

We also examined the effects of different parameters of bioinformatic assembly tools on the three *D. reticulatus* ticks and identified the best approach to obtain high-quality assemblies from deeply sequenced tick genomes. We tested the effects of several parameters on assembly, finding that Dorado corrected reads and limiting Flye to 40x coverage provided substantial improvements to the final assemblies based on the QUAST and BUSCO compleasm assembly metrics which provide the level of fragmentation and completeness of each genome. Investigating these parameters in an organism with such a large and highly repetitive genome required substantial computational resources (months of compute time on nodes with 1 TB of RAM). The clear demonstration that using Dorado corrections and modest limitations to read depth can yield substantial improvements to assemblies should provide a helpful guide to other researchers working on organisms with similarly large and repetitive genomes. Based on tick genome assembly quality parameters (Dillon et al., 2026), all three *D. reticulatus* assemblies are classified as high quality.

Chromosome number is currently unknown for *D. reticulatus*, which inhibits our understanding of this species genome. If the sequences are sufficient for large-scale assembly, then researchers face the issue of assigning chromosome-sized scaffolds to chromosomes, which is particularly difficult for organisms with multiple similar-sized chromosomes (De et al., 2023) or no published karyotype. However, our three *D. reticulatus* assemblies, the chromosome-sized *D. reticulatus* assembly (Billows et al., 2026), and our synteny graphs indicate *D. reticulatus* haploid genome has 11 chromosomes. With the availability of high-quality *Dermacentor* genome assemblies, we were able to visualize the unique and syntenic genetic regions among the species. *Dermacentor* spp. females have 22 chromosomes, and males 21 chromosomes in their diploid genomes (i.e., males are XO) (Geraci, Spencer Johnston, Paul Robinson, Wikel, & Hill, 2007; Gunn & Hilburn, 1989, 1990). Chromosome assignments are best achieved by properly anchored fluorescent *in situ* hybridization (FISH) of probes to chromosomes, but those experiments are difficult, time-consuming, and expensive. No tick genome assembly is currently chromosome-anchored (Dillon et al., 2026), but we find that orthologous gene synteny analysis allows the assignation of putative chromosome IDs for tick species within the genus *Dermacentor*. Gene synteny analysis makes use of readily available information from assemblies and does not require chromosome anchoring via FISH. It is important to note that Figure 3 emphasizes orthology of the chromosomes, but still displays scaffold numbers, which are assigned by size, but may not reflect the full size of chromosomes due to incomplete sequence data or assembly. Extending such analyses to additional taxa and types of DNA is likely to be a fruitful approach for understanding evolutionary processes and phylogenetics (Bhoi, Mule, Savale, Sharma, & Kulkarni, 2026).

We find sequencing and *de novo* assembly of multiple individuals from one species is advantageous as it allows for intra-species comparisons as well as increased confidence in inter-species comparisons. Orthogroup analyses strongly benefit from information from multiple individuals. The 920 orthogroups shared between Louise and *D. silvarum* (Figure 3), for example, would be categorized as unique to *D. silvarum* if the Louise assembly were not available. While we show that the Devon and Louise assemblies each have more than 600 unique orthogroups (Figure 3), these are likely due to differences in gene annotation approaches and gene fragmentation versus gene content variation (Cramaro et al., 2015).

We find TEs comprise at least 63% of each *D. reticulatus* assembly with manual TE curation. In comparison, using automated TE annotation, 60.76% was found for *D. reticulatus* Devon (Billows et al., 2026). We see that on average, Elska, Louise, and Penny have ∼5 times more DNA transposons, ∼26% fewer LINE elements, and ∼1.14 times more LTR elements than the Devon assembly (Billows et al., 2026). Notably, our TE curation allows more than 60% of the unknown TEs relative to the Devon assembly (Billows et al., 2026). Our manual annotation of *I. scapularis* and *I. ricinus* found 69% and 68% TEs, respectively (Ronai et al., 2026). Thus, *D. reticulatus* has fewer TEs but are LTR-rich. It should also be noted that the manually curated TEs for *I. ricinus* and *I. scapularis* do not report unknown TEs (Ronai et al., 2026). Here we report 16.5% unknown TEs that may represent uncharacterized TEs or highly degraded repetitive sequences. A benefit to manual TE curation is more accurate calculations of genetic divergence compared to homology-based methods (Goubert et al., 2022). Thus, these assemblies will be valuable to future studies that use them to calculate genetic divergence among ticks.

Variation within and among individuals was also investigated for SNPs. The number of individuals used here is well below the sampling used in population genomics of other ticks (Dong & Schoville, preprint; Frederick et al., 2023; Schoville et al., 2024), thus we limit the interpretation of our results. The samples in the Netherlands population had an average pi of 1.49% or 1.55%, depending on the reference used, whereas the UK tick had an average pi of 1.01% or 1.32%; indicating the three Netherlands individuals showed higher observed nucleotide diversity than the single UK individual, regardless of reference assembly used. All of these pi values fall within the average pi for Arthropoda (Leffler et al., 2012). The outliers observed in Figure 6 suggest areas of the genome that are worthy of additional investigation with data. Additional data are also needed to understand the population structure of *D. reticulatus*, but the two reference assemblies used here will be valuable resources for this future work.

Our study explores the impacts of a variety of choices researchers face in the amount and quality of genomic data to collect, the amount of data to use for genome assemblies, as well as the genomic complexities of ecologically relevant animals, particularly those with high TE content. We show that ONT data from R09.4.1 flow cells can and should be base called with newer versions of Guppy, then, when necessary, combined with additional data from R10.4.1 flow cells and Dorado error correction to obtain coverage limited to 40x for Flye assemblies.

## ACKNOWLEDGEMENTS

We thank Ron Dirks (Future Genomics Technologies, Leiden, The Netherlands) for invaluable advice on sample preparation and next generation sequencing. We thank Oliver Levine and Amanda Sullivan for providing expert advice in handling and modifying complex bioinformatic outputs. The Georgia Advanced Computing Resource Center provided critical resources for genome assembly and annotation. The High-Performance Computing Center at Texas Tech University provided critical resources for transposable element curation and analysis. KCD and TCG have been supported in part with federal funds from the Centers of Disease Control and Prevention Pathogen Genomics Center of Excellence (Contract No. NU50CK000626). HS was financially supported by the Dutch Ministry of Health, Welfare and Sport (VWS).

## DATA ACCESSIBILITY AND BENEFIT SHARING STATEMENTS

### Data accessibility statement

Raw reads and genome assemblies are available under the NCBI BioProject PRJNA1130541. Scripts used to generate the data and figures are located on GitHub: https://github.com/katiecdillon/Dreticulatus.

### Benefit sharing statement

A research collaboration was developed with scientists from the countries providing genetic samples, all collaborators are included as co-authors, the results of research have been shared with the provider communities, and the broader scientific community as the data and results can be found in public databases as described above.

## AUTHOR CONTRIBUTIONS

Conceptualization: KCD, HS, TCG

Validation: KCD, DAR, RPB, SC

Formal analysis: KCD, DAR, SC, CG

Investigation: KCD

Resources: HS, TCG

Data Curation: KCD, DAR

Writing - Original Draft: KCD, IR, TCG, HS, DAR, SC, CG

Writing - Review & Editing: All

Visualization: KCD, IR, DAR, SC, CG

Supervision: HS, DAR, TCG

Project administration: KCD, HS, DAR, TCG

Funding acquisition: HS

## CONFLICTS OF INTEREST

The authors declare no conflicts of interest.

## Notes

### Competing Interest Statement

The authors have declared no competing interest.

### Summary of Updates

References to Table S2, S3, and S4 were updated to be consistent with the manuscript and supplemental materials. Grammatical errors were also revised.

