## Supplementary figures and legends for "Optimizing genome assembly, chromosome synteny, and genetic variant discovery from Oxford Nanopore sequences of *Dermacentor reticulatus* ticks"

**SUPPLEMENTAL TABLE LEGENDS**

**Assembly key (S1-S5)**. From left to right, the step-by-step approach to the naming system for all unique IDs in tables S1-S6. First, the “Tick ID” for each tick specimen in the data set. Second, the ONT “Flow cell” that each tick was sequenced on. Third, “Replicate” indicates if a tick was sequenced using the same sequencing parameters more than once. Fourth, “Guppy” represents the version of the bioinformatic tool Guppy that was used for basecalling. Fifth, “Dorado” indicates whether raw sequence reads were error corrected or not with the bioinformatic tool Dorado. Sixth, “Flye assembly coverage” indicates what assembly coverage was set for each tick assembly. Seventh, “purge_haplotigs” indicates if an assembly is diploid or if the assembly was modified with the bioinformatic tool purge_haplotigs to achieve a pseudo-haploid assembly. For example, the unique ID E.1.0.1.0.0.2 represents the tick Elska, which was sequenced on an R09.4.1 flow cell, was not a replicate, basecalled with Guppy version 4.3.4, was not error corrected with Dorado, no assembly coverage parameter set, and the assembly is diploid.

**Table S1. Nanoplot assessment for 19 raw tick sequences.** For the raw tick sequence reads were assessed raw read quality metrics using Nanoplot version 1.33.0 and version 1.42.0. Each raw tick sequence read has a unique ID (see key). The mean and median read length and read quality is reported. The number of reads, read length N50, standard deviation read length, and total bases are reported. The number of bases at Phred quality score cutoffs greater than Q5, Q7, Q10, Q12, Q15 are reported for raw sequence reads prior to Dorado correct. The Dorado correct output is in FASTA format and does not report quality scores.

**Table S2. QUAST assessment for 38 tick genome assemblies.** For the tick genome assemblies to assess assembly quality metrics QUAST-LG (v5.2.0) was used. Each tick assembly has a unique ID, following the key. The number of contigs and the total length of each assembly is reported (>= 0 bp, >= 1,000 bp, >= 3,000 bp, >= 5,000 bp, >= 10,000 bp, >= 25,000 bp, >= 50,000 bp). The length of the largest contig is provided. Total length is the total number of bases, including Ns. GC% represents the proportion of G and C nucleotides in the assembly relative to its total length. N50 and N90 are the lengths in which all contigs equal to or longer than that length account for at least 50% or 90% of the total assembly, respectively. auN (area under the N curve), is a more comprehensive statistic than N50, as it is less affected by contig length and considers the entire Nx curve. L50 and L90 represent the smallest number of contigs that cover 50% or 90% of the total assembly, respectively. The number of Ns per 100 kbp represents uncalled and unmasked bases in the assembly.

**Table S3. Raw reads before and after Dorado read error correction.** Raw read calculations for combined raw reads before and after Dorado read error correction. Unique ID, total bases, and mean read length are copied from Table S1. The percent of total bases retained after Dorado read error correction was calculated. The total base coverage is calculated from the average QUAST total length of the final high-quality assemblies for Elska, Louise, and Penny. The mean read length difference is calculated from subtracting the mean read length post-Dorado from the mean read length pre-Dorado.

**Table S4. BUSCO assessment for 38 tick genome assemblies.** For the tick genome assemblies to assess assembly completeness metrics with compleasm (v0.2.5) was used. Each tick assembly has a unique ID, following the key. Fragmented genes of subclass 1 and subclass 2 can be partially aligned to the assembly where one portion can or cannot be aligned at another position, respectively. Missing genes are unable to be aligned to the assembly.

**Table S5. Bacterial contaminants identified in the highest quality tick assemblies for each individual.** Contaminants within each pseudo-haploid assembly for Elska, Louise, and Penny were identified and removed using the NCBI Foreign Contamination Screen (v0.5.0) genome cross-species (FCS-GX) tool. For FCS-GX, tax-id 34619 was used. Additional bacterial contaminants were found and removed from the final, soft-masked Penny assembly after submission to NCBI GenBank. For each bacterial contaminant, the contig it was found on is listed, the location on the contig, the sequence length of the contig, the FCS-GX recommendation for treatment, taxonomic division, aggregate contig coverage, and the taxonomic name for the bacterial contaminant. Additionally, we calculate the length of each contaminant based on the start and stop position on the contig. If a genome assembly of the contaminant was available on NCBI RefSeq, we list the assembly length, checkM completeness, and the RefSeq URL to the genome assembly.

**Table S6. Adaptor contaminants identified in the highest quality tick assemblies for each individual.** Contaminants within each pseudo-haploid assembly for Elska, Louise, and Penny were identified and removed using the NCBI Foreign Contamination Screen (v0.5.0) adaptor (FCS-adaptor) tool. For FCS-adaptor, the ‘-euk’ tag was used. For each adaptor contaminant, the contig it was found on is listed, the sequence length of the contig, the FCS-GX recommendation for treatment, the range is the coordinate span of the contig where the contaminant is found, and the name of the contaminant.

**Table S7. High-quality tick assemblies for each individual tick (Elska, Louise, and Penny).** Each assembly was uploaded to NCBI GenBank where additional bacterial contaminants were identified and removed from the Penny assembly. Assembly quality metrics QUAST-LG (v5.2.0) and completeness metrics with BUSCO compleasm (v0.2.8) were assessed for each assembly. The NCBI GenBank ID is listed for each assembly representing a single tick. The number of contigs (>= 0 bp) and the total length of each assembly is reported. Total length is the total number of bases, including Ns. GC% represents the proportion of G and C nucleotides in the assembly relative to its total length. N50 and N90 are the lengths in which all contigs equal to or longer than that length account for at least 50% or 90% of the total assembly, respectively. auN (area under the N curve), is a more comprehensive statistic than N50, as it is less affected by contig length and considers the entire Nx curve. L50 and L90 represent the smallest number of contigs that cover 50% or 90% of the total assembly, respectively. The number of Ns per 100 kbp represents uncalled and unmasked bases in the assembly. A total of 1,667 BUSCO genes from the phylum Arthropoda (arthropoda_odb12) were used to assess each assembly.

**Table S8. Proportion of repetitive elements in *D. reticulatus*.** Proportion of repeats in the Elska, Louise, and Penny *D. reticulatus* pseudo-haploid assemblies. The data was obtained from the RepeatMasker .out files for each tick, and each repetitive element is separated by class, subclass, number of base pairs, and proportion of each genome assembly.

**Table S9. Population genetics summary statistics of *D. reticulatus* assemblies.** Population genetics summary statistics were calculated for each chromosome using the bioinformatic tool pixy (v2.0.0) following the methods outlined here: <https://github.com/katiecdillon/Dreticulatus/tree/main/pixy_plots>. During the alignment stage, the *D. reticulatus* Louise assembly was used as reference and the contigs that mapped to the UK assembly’s eleven chromosome-sized scaffolds were used. During the alignment stage, the *D. reticulatus* UK assembly was used as reference. The “Netherlands” population consists of three samples: Elska, Louise, and Penny. The “UK” population consists of one sample: UK. The lowest and highest values are highlighted in green and red, respectively. **A)** Pi and Watterson’s theta summary statistics were calculated for each chromosome-sized scaffold and split between the two populations. **B)** The d_xy_ and Hudson’s F_ST_ summary statistics were calculated for each chromosome-sized scaffold for both populations combined. **C)** The Tajima’s D summary statistic was calculated for each chromosome-sized scaffold for the Netherlands population only.

**SUPPLEMENTAL FIGURES**

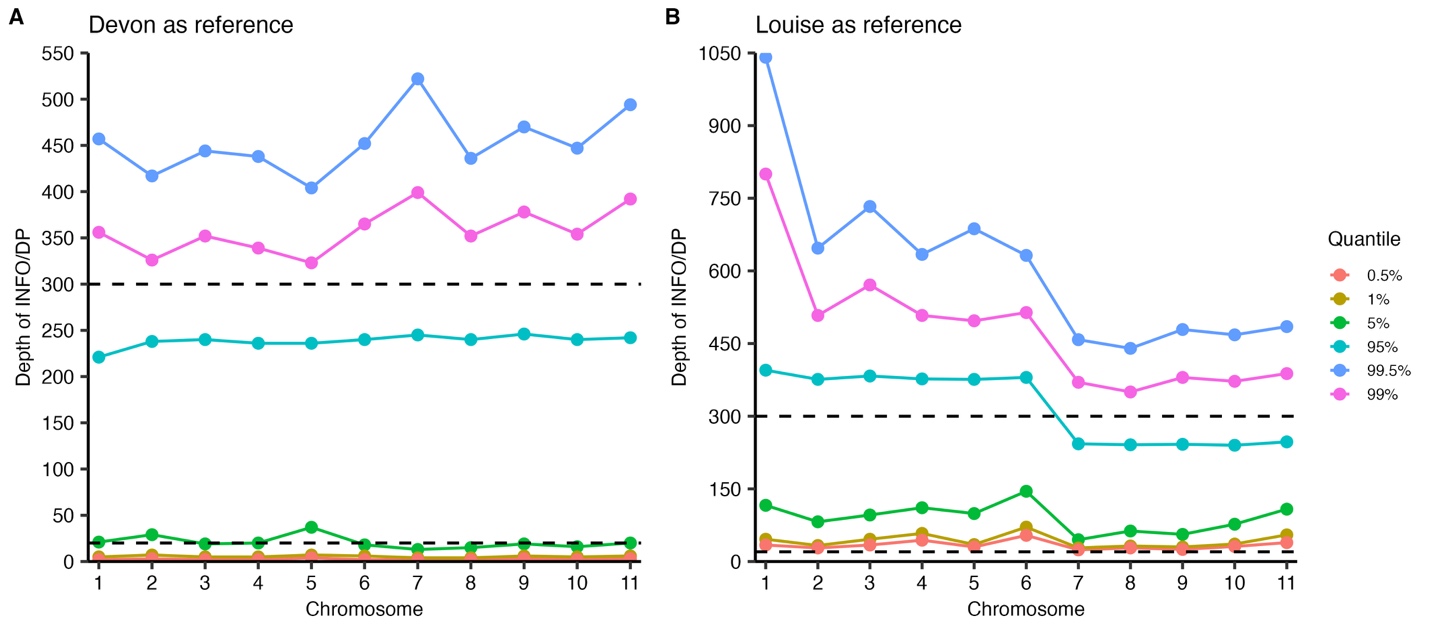

**Figure S1.** **Depth thresholds for VCF filtering.** After variants were called and output to a VCF format, each VCF file required quality filtering prior to running pixy. Each colored line represents a quantile in which the values in the INFO/DP column of each VCF file fell into. The dotted lines represent the thresholds between 20x-300x coverage. **A)** The *D. reticulatus* Devon assembly was used as the reference during the alignment stage. The y-axis goes up to 550x. **B)** The *D. reticulatus* Louise assembly was used as the reference during the alignment stage. The y-axis goes up to 1,050x.

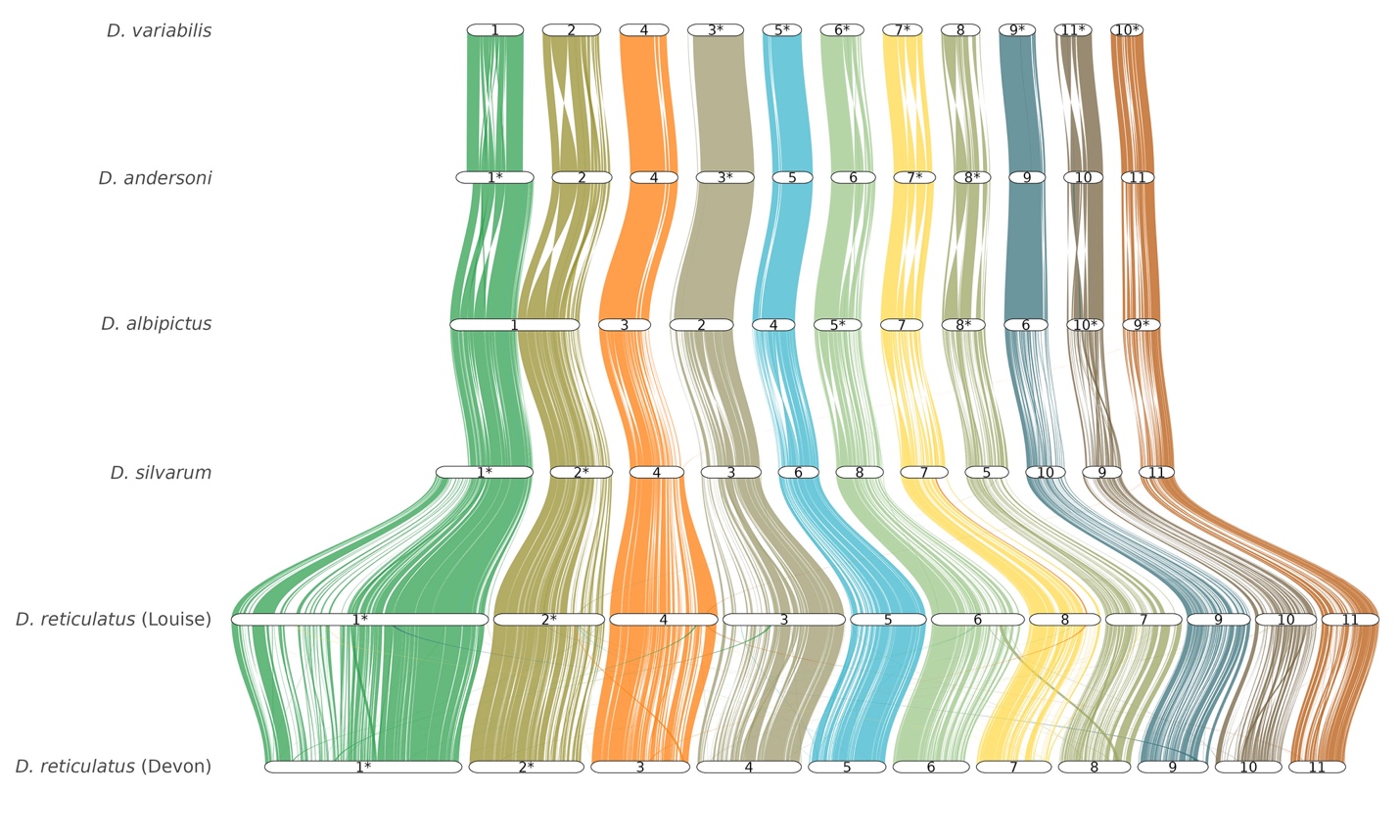

**Figure S2. Synteny of the longest isoform transcripts in chromosome-sized scaffolds in *Dermacentor* spp. ticks**. Tick assemblies are ordered according to the OrthoFinder species tree starting with *D. reticulatus* (Devon). Asterisks (*) note the chromosome-sized scaffolds that were manually flipped in GeneSpace. The longest isoforms per gene were extracted and used to generate syntenic blocks. Chromosome-sized scaffolds scaled by gene rank order.

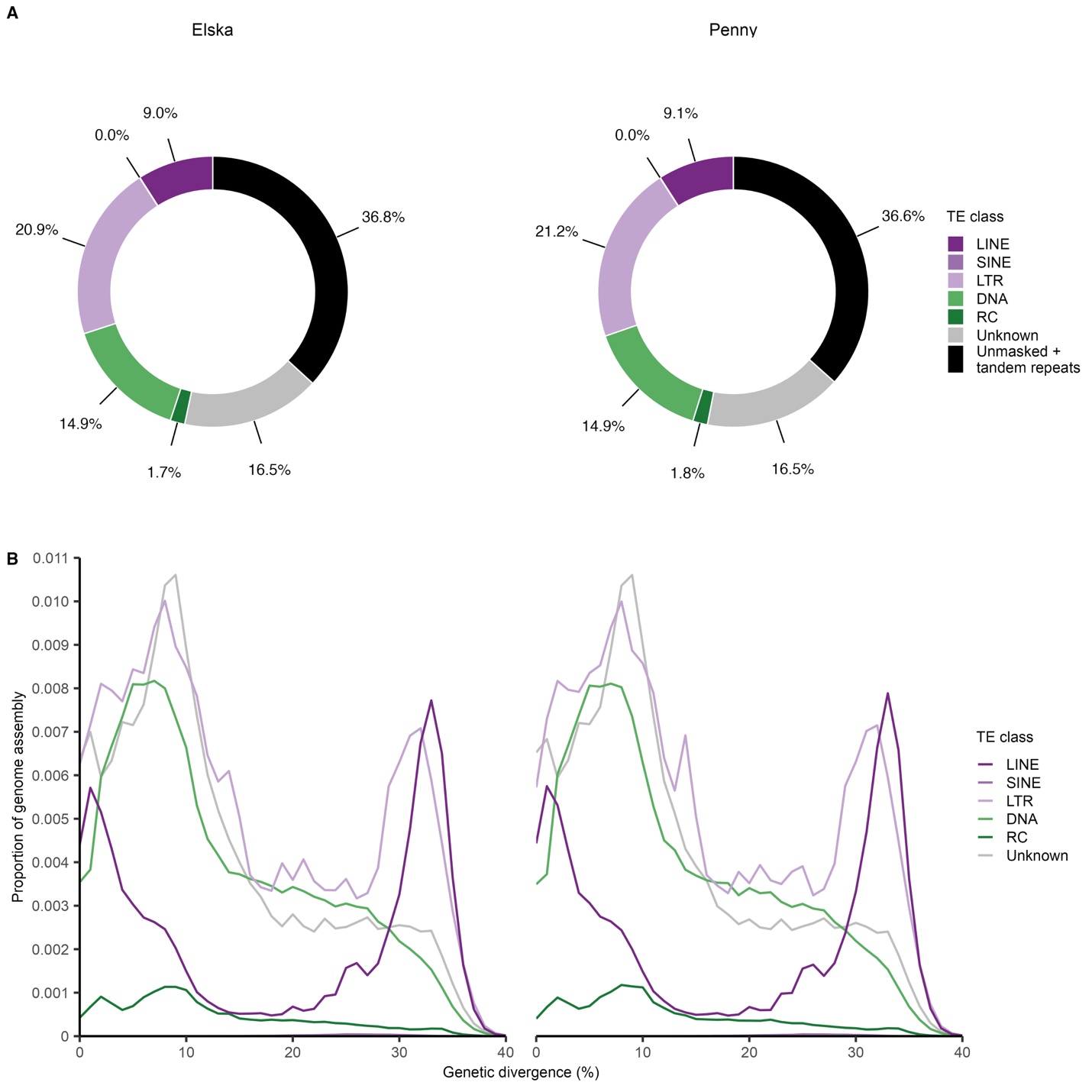

**Figure S3. Characterization of the transposable elements in *D*. *reticulatus* Elska and Penny.** The transposable element landscape of *D. reticulatus* Elska and Penny. **A)** Proportion of each assembly occupied by the major TE classes, unknown TEs, unmasked sequences, and tandem repeats. **B)** Repetitive proportion of TE classes and genetic divergence in each assembly.

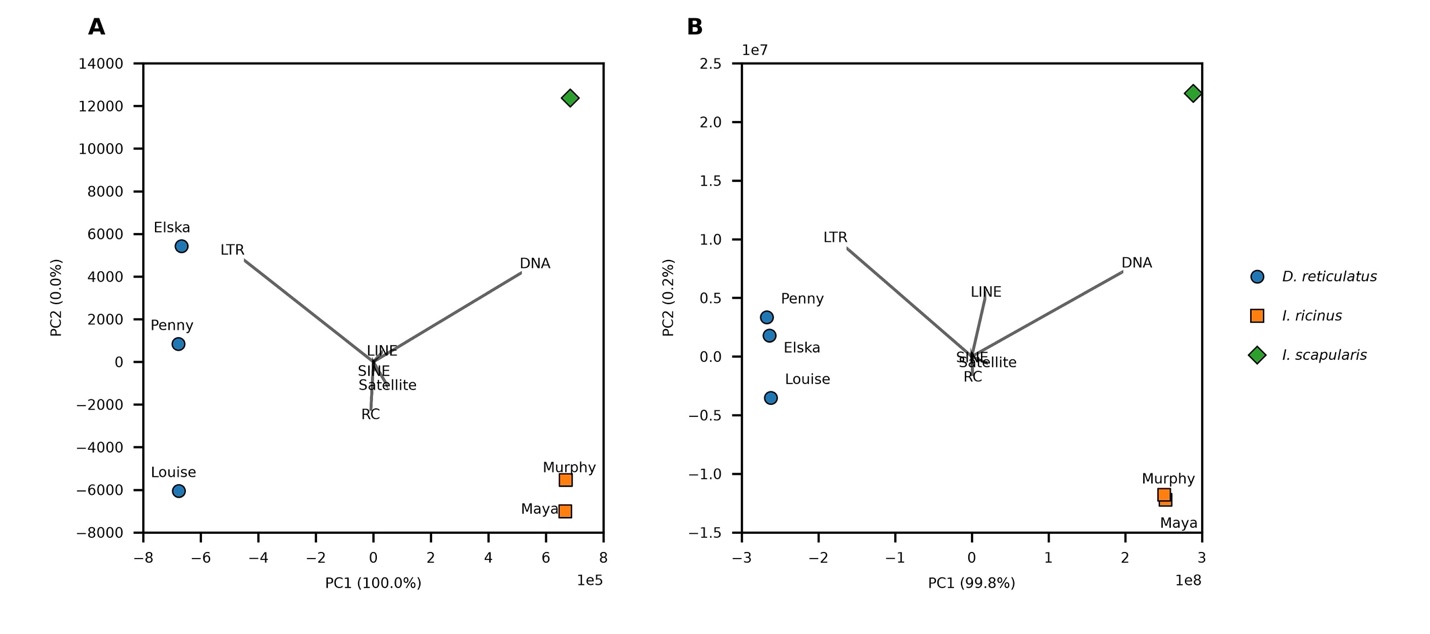

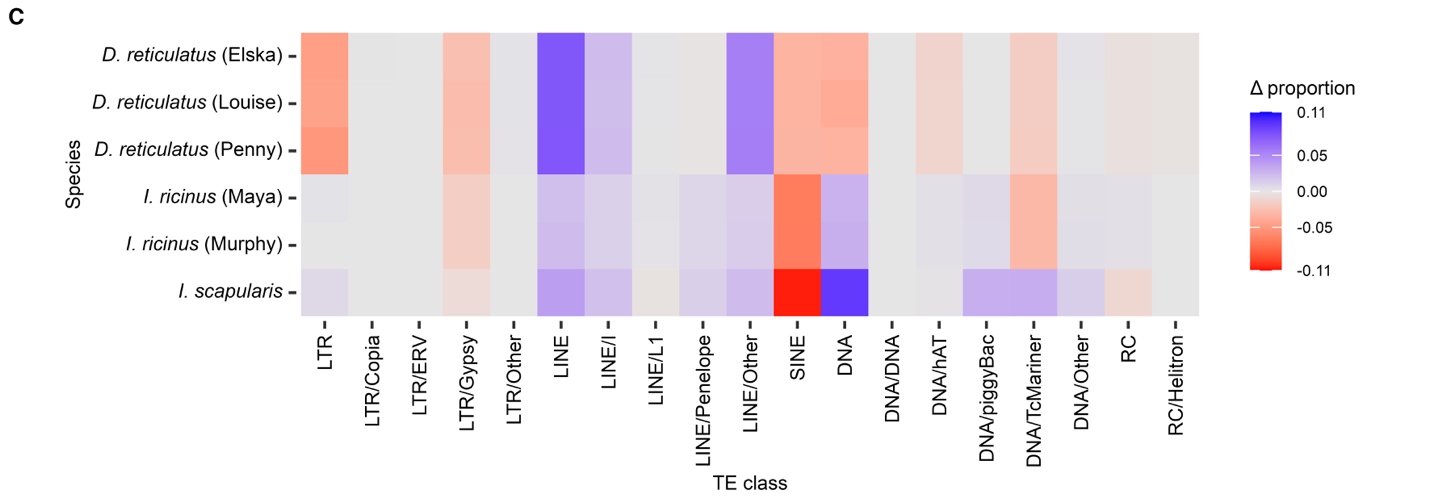
**Figure S4. Comparison of TE classes in the six tick genome assemblies with manually curated TEs.** The three *D. reticulatus* assemblies (this paper), *I. ricinus* Maya (GCA_043698115.1) and Murphy (GCA_043645445.1), and *I. scapularis* (GCA_016920785.2). **A)** Principal Component Analyses biplot of TE occupancy to compare overall trends of TE accumulation and activity performed for each TE class using both overall proportion in representative genome assemblies of *D. reticulatus*, *I. ricinus*, and *I. scapularis*. All information is on axis one. **B)** Principal Component Analyses biplot of TE insertion counts to compare overall trends of TE accumulation and activity performed for each TE class using individual TE insertion counts. The majority of information is on axis one. **C)** Comparison of TE accumulation divergence of the six tick assemblies. Heatmap illustrating differential TE accumulation recently (divergences from the consensus < 10%) and in the more distant past (divergences from the consensus > -11%).

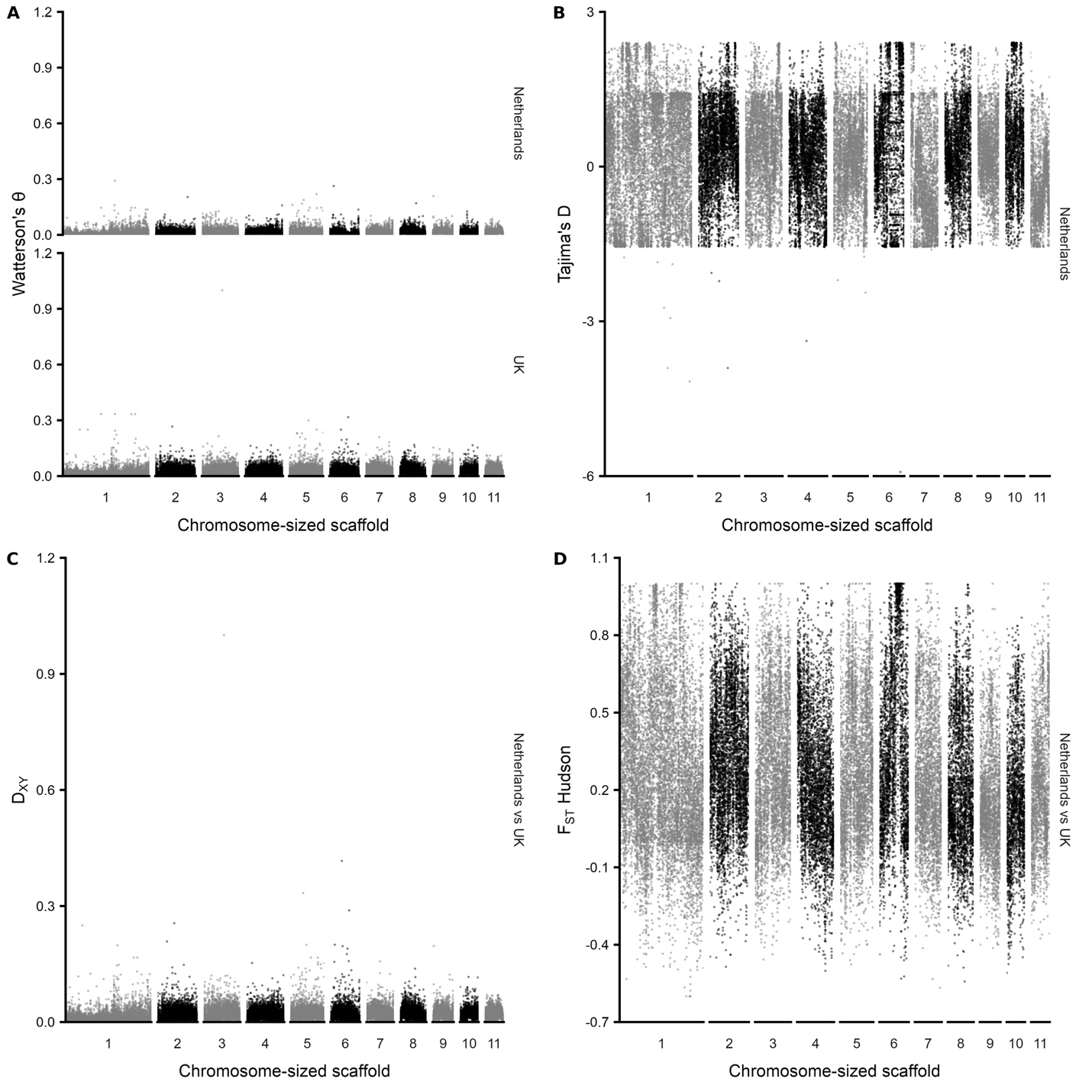

**Figure S5. Per chromosome population summary statistics where Louise is the reference.** Each plot represents a population genetics summary statistic generated by the bioinformatic tool pixy. During the alignment stage, the *D. reticulatus* Louise assembly was used as the reference and the contigs that mapped to the UK assembly’s eleven chromosome-sized scaffolds were used. The “Netherlands” population consists of three samples: Elska, Louise, and Penny. The “UK” population consists of one sample: UK.  **A)** The Watterson’s theta summary statistic calculated for each chromosome-sized scaffold and split between the two populations. **B)** The Tajima’s D summary statistic calculated for each chromosome-sized scaffold for the Netherlands population only. **C)** The d_xy_ summary statistic calculated for each chromosome-sized scaffold for both populations combined. **D)** The Hudson’s F_ST_ summary statistic calculated for each chromosome-sized scaffold for both populations combined.

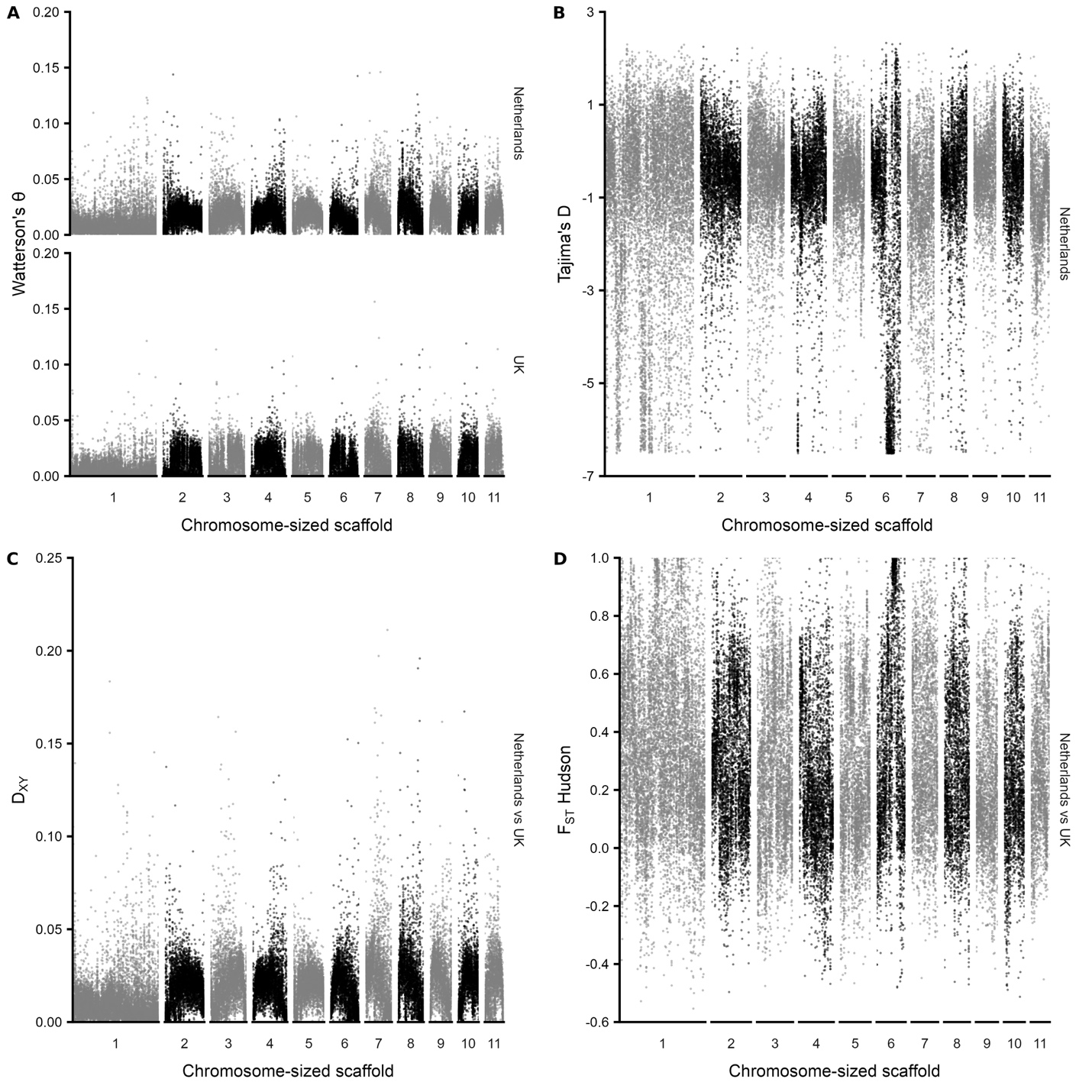

**Figure S6. Per chromosome population summary statistics where Devon is the reference.** Each plot represents a population genetics summary statistic generated by the bioinformatic tool pixy. During the alignment stage, the *D. reticulatus* UK assembly was used as reference. The “Netherlands” population consists of three samples: Elska, Louise, and Penny. The “UK” population consists of one sample: UK.  **A)** The Watterson’s theta summary statistic calculated for each chromosome-sized scaffold and split between the two populations. **B)** The Tajima’s D summary statistic calculated for each chromosome-sized scaffold for the Netherlands population only. **C)** The d_xy_ summary statistic calculated for each chromosome-sized scaffold for both populations combined. **D)** The Hudson’s F_ST_ summary statistic calculated for each chromosome-sized scaffold for both populations combined.

**SUPPLEMENTAL FILE LEGENDS**

**File S1. BUSCO of the BRAKER3 Augustus proteins in the Louise assembly.** Genes were predicted in the *D. reticulatus* Louise assembly using the BRAKER3 pipeline. The AUGUSTUS protein amino acid FASTA was assessed using compleasm. Each BUSCO ID detected in the FASTA included the BUSCO status (single, duplicate, fragmented, missing), amino acid sequence the BUSCO was found on, BUSCO score, and length of the BUSCO.

**File S2. Unique and shared orthogroups among three tick genome assemblies.** The number of orthologous gene groups identified by OrthoFinder that are unique to, or shared among, *D. reticulatus* (Devon guided assembly), *D. reticulatus* (Louise assembly), and *D. silvarum* (BIME_Dsil_1.4 assembly).
